# Pannexin 1 Channels Control Cardiomyocyte Metabolism and Neutrophil Recruitment During Non-Ischemic Heart Failure

**DOI:** 10.1101/2023.12.29.573679

**Authors:** Caitlin M. Pavelec, Alexander P. Young, Hannah L. Luviano, Emily E. Orrell, Anna Szagdaj, Nabin Poudel, Abigail G. Wolpe, Samantha H. Thomas, Scott Yeudall, Clint M. Upchurch, Mark D. Okusa, Brant E. Isakson, Matthew J. Wolf, Norbert Leitinger

## Abstract

Pannexin 1 (PANX1), a ubiquitously expressed ATP release membrane channel, has been shown to play a role in inflammation, blood pressure regulation, and myocardial infarction. However, a possible role of PANX1 in cardiomyocytes in the progression of heart failure has not yet been investigated. We generated a novel mouse line with constitutive deletion of PANX1 in cardiomyocytes (Panx1^MyHC6^). PANX1 deletion in cardiomyocytes had no effect on unstressed heart function but increased the glycolytic metabolism both *in vivo* and *in vitro*. *In vitro*, treatment of H9c2 cardiomyocytes with isoproterenol led to PANX1-dependent release of ATP and Yo-Pro-1 uptake, as assessed by pharmacological blockade with spironolactone and siRNA-mediated knock-down of PANX1. To investigate non-ischemic heart failure and the preceding cardiac hypertrophy we administered isoproterenol, and we demonstrate that Panx1^MyHC6^ mice were protected from systolic and diastolic left ventricle volume increases and cardiomyocyte hypertrophy. Moreover, we found that Panx1^MyHC6^ mice showed decreased isoproterenol-induced recruitment of immune cells (CD45^+^), particularly neutrophils (CD11b^+^, Ly6g^+^), to the myocardium. Together these data demonstrate that PANX1 deficiency in cardiomyocytes impacts glycolytic metabolism and protects against cardiac hypertrophy in non-ischemic heart failure at least in part by reducing immune cell recruitment. Our study implies PANX1 channel inhibition as a therapeutic approach to ameliorate cardiac dysfunction in heart failure patients.

## Introduction

Heart failure, which affects five million Americans each year and has a five-year survival rate of less than 50 percent^1^, can be divided in to two sub-pathologies: heart failure with reduced ejection fraction (HFrEF) and heart failure with preserved ejection fraction (HFpEF); HFpEF is the more predominant form and its prevalence is on the rise^1,2^. While several effective therapies are available for patients with HFrEF, HFpEF has limited treatment options underscoring the need to identify new therapeutic strategies^3^. Pathological features of heart failure include cardiac hypertrophy, chronic inflammation, and interstitial fibrosis of the heart muscle, which together result in reduced cardiac contractile function^4^. Current pharmacologic interventions for HFrEF include treatments with beta-blockers, loop diuretics, angiotensin converting enzyme (ACE) inhibitors, angiotensin receptor blockers, and sodium glucose transporter 2 (SGLT2) inhibitors^4–6^. These therapies target the renin-angiotensin/ aldosterone systems and the adrenergic signaling axes, providing more symptom management than improving function of the cardiac muscle^5^. Moreover, previous clinical trials that have targeted TNF-α signaling, modulated the immune system using intravenous immunoglobulins^7^, or decreased fibroblast activation with pirfenidone^8^ (an antifibrotic which down regulates growth factors and procollagens I and II), have not lead to significant improvements in patient outcomes. To prevent cardiac muscle decline, it is necessary to devise novel therapeutic approaches which directly target the mechanisms of heart failure progression.

Proper contractile function of the heart requires, at the level of the cardiomyocyte, sufficient cellular energy in the form of accessible ATP^9,10^. Adult cardiomyocytes rely on fatty acids as their main fuel source to generate ATP, but they are metabolically flexible and can use glucose, branched-chain amino acids, or ketones as needed^11^. During heart failure, cardiomyocytes demonstrate an increased demand for ATP due to more frequent but less effective contraction^12^. In order to meet this increased energy demand, cardiomyocytes shift their metabolism from utilizing fatty acids to utilizing glucose, which is accompanied by increased glycolysis^13^, which resembles the reliance on glucose and glycolysis in fetal cardiomyocytes^11,14^. Moreover, recent work has demonstrated that decreased reliance on fatty acid oxidation confers greater resistance to ischemia reperfusion injury and promotes cardiomyocyte proliferation in mice deficient in the mitochondrial fatty acid transporter *Cpt1b*^15^. These findings suggest that a metabolic shift to glucose utilization by cardiomyocytes protects against cardiac injury.

In response to local hypoxia or chronic beta-adrenergic stimulation, two conditions known to cause cardiac hypertrophy and heart failure, cardiomyocytes release damage signals such as ATP^16^ and secrete pro-inflammatory cytokines, which leads to the recruitment of macrophages and neutrophils and efferocytosis of dying cardiomyocytes^7,17^. Initially, recruited leukocytes promote the repair of damaged myocardium by stimulating resident fibroblasts to secrete extracellular matrix, and by inducing angiogenesis to provide needed oxygenation and nutrients to the healing cardiac muscle^18,19^. While this acute inflammatory response is necessary to activate fibroblasts and prevent excessive cardiomyocyte death, chronic activation of both the innate and adaptive immune responses has been shown to exacerbate heart failure progression^7,20–22^. The immune cell cascade and timeline for resolution of inflammation after ischemic myocardial infarction has been well characterized, however, it is less well understood how immune cells are recruited to the injured myocardium in chronically developing non-ischemic heart failure^23^.

Pannexin 1 (PANX1) is a ubiquitously expressed transmembrane channel, through which stressed or damaged cells release ATP and other small metabolites that are smaller than 1 kDa. PANX1-dependent ATP release from apoptotic cells serves as a “find me” signal for efferocytosis by immune cells^24–26^. PANX1 channel opening can be activated via multiple mechanisms including α_1D_^27^- or β-adrenergic receptor stimulation^28,29^, and caspase-3-mediated cleavage^30^. Several pharmacologic compounds including probenecid, carbenoxolone, and the FDA-approved heart failure therapeutic spironolactone have been identified as PANX1 channel blockers^31–33^. Release of ATP from cardiomyocytes through PANX1, which caused fibroblast activation *in vitro,* has previously been described^16^. Moreover, global PANX1 deletion in mice decreased ischemic area in a model of cardiac ischemia reperfusion injury^34^, while endothelial cell-specific deletion of PANX1 in mice decreased immune cell infiltration in a model of myocardial infarction^35^. Though much of the work done with PANX1 heavily implicates the role for PANX1 dependent ATP release, previous studies have demonstrated that a wide variety of metabolites can be released through PANX1. Other metabolites including ADP, UTP, and UDP-glucose have been identified in the supernatant of media collected from PANX1 dependent activation^25^. These metabolites serve as ligands for many purinergic receptors present on leukocytes, including P2Y14, P2Y2, and P2X1 which have been previously demonstrated to play key roles in chemotaxis and leukocyte migration into tissues during disease^36–39^ . While these prior studies have strongly implied PANX1 as a therapeutic target for cardiovascular diseases in general^40^, the role for PANX1 in cardiomyocytes specifically in the regulation of chronic cardiac pathologies, including heart failure, remains to be elucidated.

Here we report that PANX1 channels can be activated in cardiomyocytes through beta-adrenergic receptor stimulation. Using mice with a cardiomyocyte-specific PANX1 deletion (Panx1^MyHC6^), we demonstrate a crucial role for PANX1 channels in the development of non-ischemic heart failure. Panx1^MyHC6^ mice were protected against isoproterenol-induced ^41,42^ cardiac hypertrophy, systolic dysfunction, and neutrophil infiltration into the myocardium. Furthermore, we find that PANX1 deletion facilitates glucose utilization and increases glycolytic and decreases inflammatory gene expression. Taken together our findings implicate PANX1 channels as promising therapeutic targets in non-ischemic heart failure and cardiac hypertrophy.

## Results

### Isoproterenol Activates Pannexin 1 channels in Cardiomyocytes

Previous studies have demonstrated that Pannexin 1 (PANX1) channels are activated in response to β3 and α_1D_-adrenergic receptor stimuli^27,28^. A recent study demonstrates beta-adrenergic receptor 1-dependent activation of Panx1 channels in cardiomyocytes via cAMP/PKA-mediated phosphorylation on Serine 206^29^. To activate Panx1 channels we used isoproterenol, a pan β-adrenergic receptor agonist, in H9c2 rat myoblasts which endogenously express all β-adrenergic receptor family members and Panx1^43–45^. We tested Panx1 channel opening by measuring both ATP release into the culture media and dye (Yo-Pro-1) uptake into cells. After stimulation with 2 or 20μM isoproterenol, H9c2 cells released significantly more ATP into the extracellular media compared to vehicle stimulation (**Figure 1A**). Isoproterenol-induced ATP release was abolished by either siRNA mediated knock-down of Panx1 (successful knock-down is demonstrated by significantly decreased mRNA and protein levels (S1A,B)), or pharmacological channel inhibition with spironolactone^33^ (**Figure 1A,C**). Cell viability was not affected at either dose of isoproterenol (S1C).

**Figure 1:**
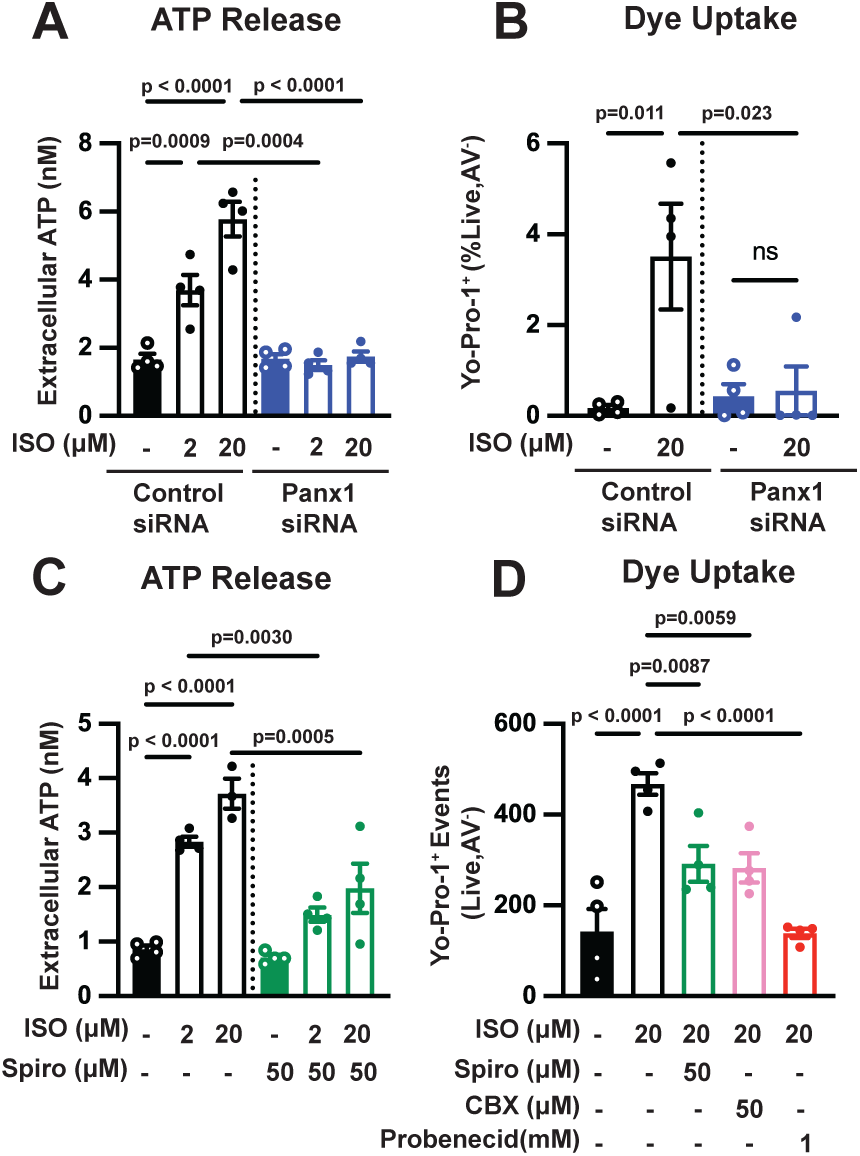
Beta-adrenergic receptor stimulation activates Pannexin 1 channels in cardiomyocytes *in vitro* A) H9c2 were transfected with either control or Panx1-targeting siRNA for 72 hours and extracellular concentration of ATP was measured after 15 minutes of stimulation with isoproterenol (ISO) (2 and 20μM) or vehicle control. (N=4) B) Flow cytometric analysis of Yo-Pro-1 dye uptake by control of Panx1-targeting siRNA treated H9c2 cells after stimulation with 20μM ISO (gated for live, annexin V^-^ cells). (N=4) C) H9c2 cells were pretreated with spironolactone (50μM) or vehicle control and extracellular ATP levels were measured after 15 minutes of stimulation with isoproterenol (ISO) (2 and 20μM). (N=4) D) Yo-Pro-1 dye uptake by H9c2 cells treated with isoproterenol (20μM) after pretreatment with spironolactone, carbenoxolone (CBX), or probenecid at the listed concentrations. (N=4) Data are represented as mean ± standard error of the mean (SEM). Data shown are representatives of 2 independent experiments. Significance determined using One-way ANOVA (A, B, C, D) with Sidak’s multiple comparison test for post-hoc analysis for comparisons between individual groups. ns = not significant See also S1 in reference to figure 1.

Additionally, using a flow-cytometry based approach, we found that stimulation with 20μM isoproterenol caused a significant increase in Yo-Pro-1 positive, Annexin V negative H9c2 cells compared with vehicle stimulation (**Figure 1B**, representative gating S1D). Isoproterenol-induced Yo-Pro-1 uptake was significantly decreased after siRNA-mediated Panx1 knock-down (**Figure 1B**, S1D), and blunted with PANX1 channel blockade by spironolactone, carbenoxolone, or probenecid (**Figure 1D**). Taken together, these data demonstrate that PANX1 channel opening in cardiac myoblasts as a result of β-adrenergic receptor stimulation leads to the release of ATP.

### Generation and characterization of a mouse with Pannexin 1 deletion specifically in cardiomyocytes

To investigate the role of PANX1 channels in cardiomyocytes *in vivo*, we generated a cardiomyocyte-specific PANX1 null mouse by crossing Panx1 floxed mice^46^ with mice expressing Cre recombinase under the control of the alpha myosin heavy chain promoter (MyHC6). Confirmation of successful recombination was demonstrated by the presence of a null band on PCR analysis of genomic DNA isolated from the hearts of Panx1^MyHC6^ mice, which was not present in the hearts of Panx1^fl/fl^ mice or other tested organs (liver, kidney) from either mouse genotype (S2A). Levels of *Panx1* mRNA (by qPCR) and PANX1 protein (by immunofluorescence) were significantly decreased in isolated cardiomyocytes and cardiac troponin t (cTnnt) positive cells, respectively, from Panx1^MyHC6^ mice (**Figure 2A**, S2B) demonstrating the specificity of the Panx1 deletion in cardiomyocytes. Additionally, we see no concomitant changes in *Panx2*, *Panx3*, *Gja1* (Cx43), or *Gja5* (Cx40) mRNA (by qPCR) in isolated cardiomyocytes from Panx1^MyHC6^ mice compared with Panx1^fl/fl^ mice (S2C). Panx1^MyHC6^ mice had no obvious gross developmental defects when compared to their Panx1^fl/fl^ littermates as examined by body mass, organ mass, or tibial length at 12 weeks of age (S2D-G). Furthermore, assessment of cardiac function via 2D- echocardiography showed no significant differences in end-systolic or end-diastolic volume, wall thickness, or ejection fraction at baseline (**Figure 2B-E**, S2L). Finally, blood pressure measurements by radiotelemetry in 12-week-old male mice demonstrated no differences in mean arterial pressure, systolic or diastolic pressure, or heart rate in Panx1^MyHC6^ mice when compared to littermate Panx1^fl/fl^ mice (S2H-K).

**Figure 2:**
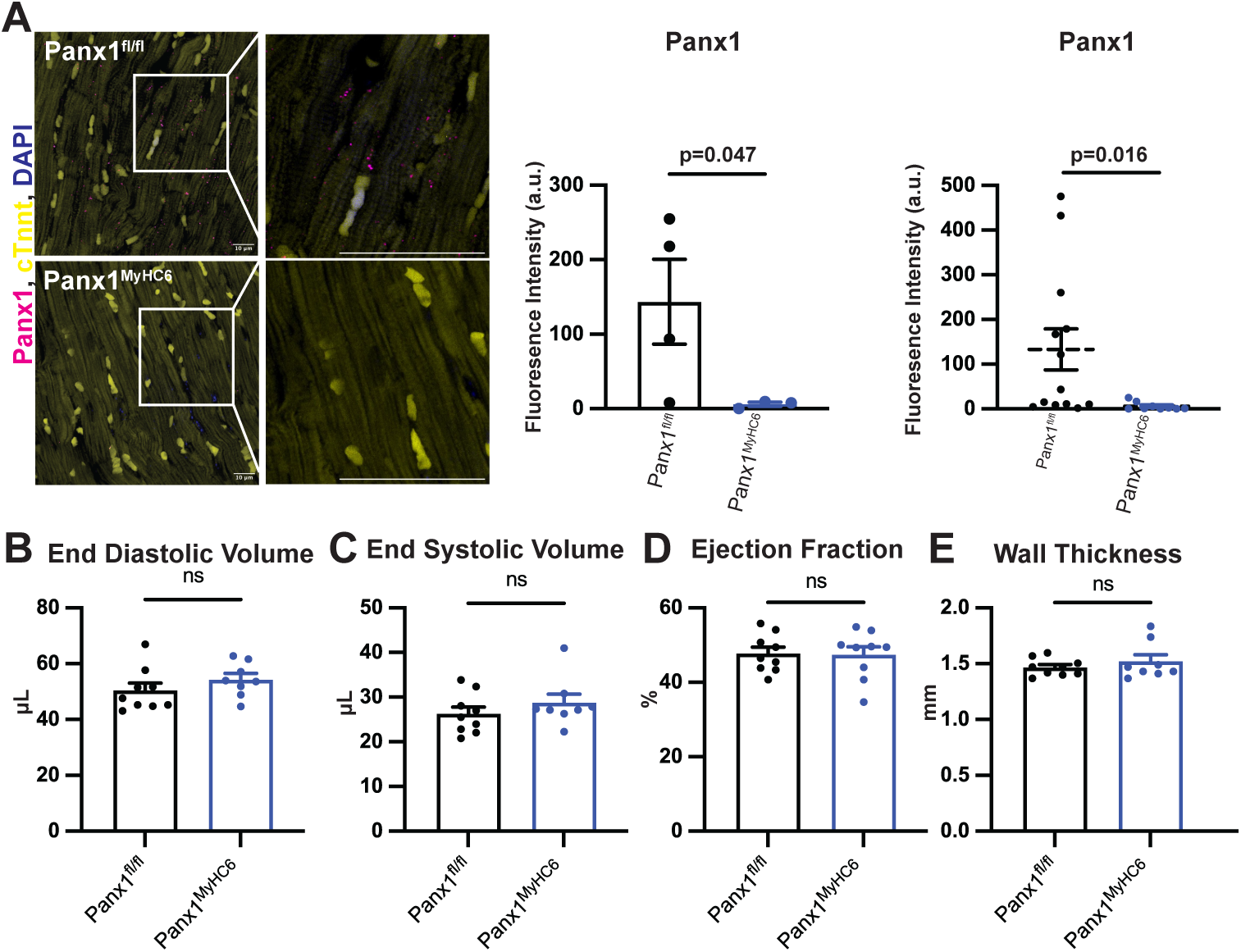
Generation and Characterization of a novel Pannexin 1 cardiomyocyte specific deletion mouse. A) Confocal micrographs of cardiomyocyte-rich whole heart sections from 12-week-old male Panx1^fl/fl^ and Panx1^MyHC6^ mice. Knockout of PANX1 in cardiomyocytes was assessed using immunofluorescent staining. Cardiac troponin T (cTnnt) was used to stain cardiomyocytes; DAPI was used as a nuclear stain. Left, representative micrographs with insets showing PANX1 punctate; middle, average Panx1 fluorescence intensity for individual mice; right, fluorescence intensities for individual micrographs showing demonstrating of PANX1 signal. (Panx1^fl/fl^ N=4 mice N=13 images, Panx1^MyHC6^ N=3 mice N=9 images, 3-5 images per mouse averaged). B) End-diastolic volume C) End-systolic volume D) Ejection fraction and E) Wall thickness were measured in anesthetized 8-to 10-week-old male Panx1^fl/fl^ and Panx1^MyHC6^ mice using *in vivo* 2D-echocardiography. (Panx1^fl/fl^ N= 9 mice, Panx1^MyHC6^ N=8) Data represented as mean ± SEM. Significance was determined by Welch’s t-test (A-E). ns = not significant. See also S2 in reference to figure 2.

### Pannexin 1 deletion shifts cardiomyocyte metabolism from oxidative phosphorylation to glycolysis

To explore effects of PANX1 deficiency on the cardiomyocyte transcriptome *in vivo*, we performed bulk RNA-sequencing on cardiomyocytes isolated from hearts of Panx1^MyHC6^ and Panx1^fl/fl^ mice, which identified 1281 and 727 uniquely expressed genes, respectively (**Figure 3A)**. There were 1069 significantly differentially expressed genes between the genotypes, with 793 down-regulated and 276 up-regulated (**Figure 3B**). Gene ontology (GO) Term and KEGG pathway analysis revealed that Panx1 deletion mainly affected pathways involved in intercellular communication, cell adhesion and gap junction function, namely “cell-cell” and “Wnt signaling”, “microtubule-based movement”, “ECM-receptor interactions”, “leukocyte trans endothelial migration”, and “cell adhesion molecules”, among others involved in inflammation and cytoskeletal organization (S3A, B). KEGG Pathway analysis was corroborated by significantly lower expression of transcripts of the claudin and cadherin family and their receptors (*Cdh1, Cldn7, Cdhr3, Cdhr4*), as well as increased expression of Transport and Golgi Organization protein 2 Homolog (*Tango2*) , which has been associated with altered mitochondrial energy metabolism in humans^47,48^ (**Figure 3B)**.

**Figure 3:**
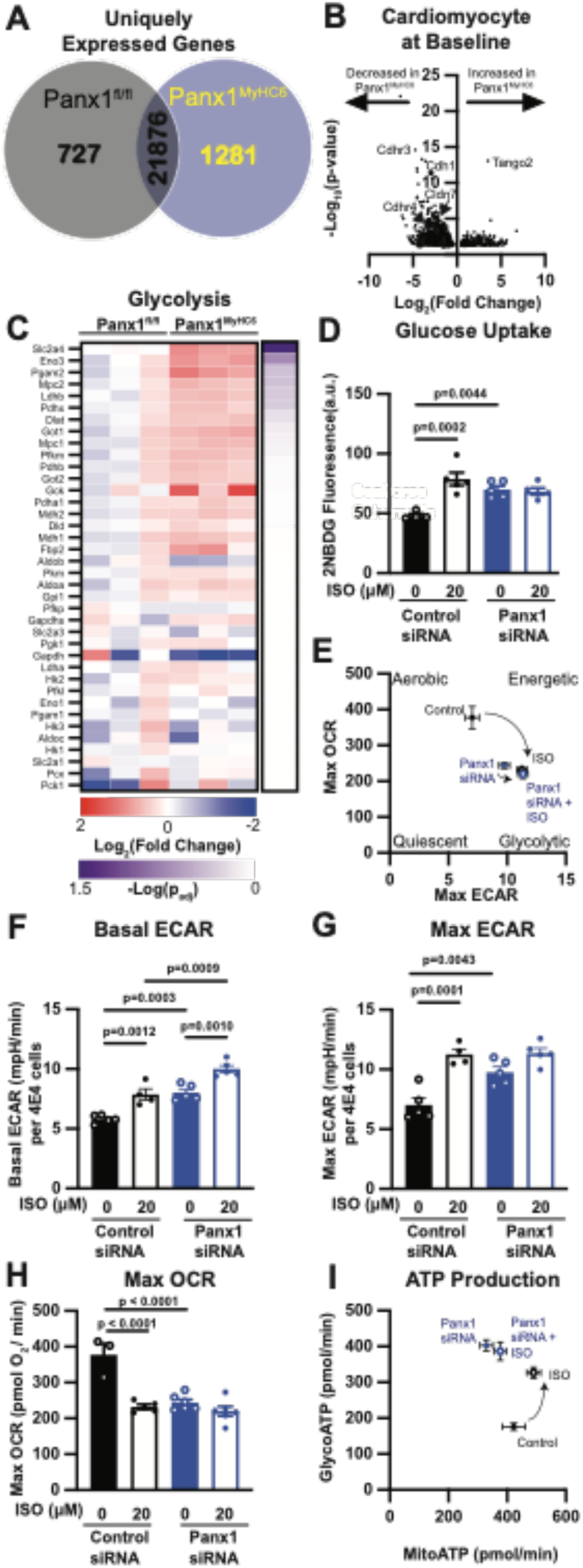
Pannexin 1 deletion shifts cardiomyocyte metabolism from oxidative phosphorylation to glycolysis both *in vivo* and *in vitro* A) Quantification of uniquely expressed genes from bulk RNA-Sequencing of isolated cardiomyocytes from untreated 12-week-old male Panx1^fl/fl^ and Panx1^MyHC6^ mice. B) Volcano plot of differentially expressed genes with a cut off of p < 0.05. Individual genes are indicated with gene name and arrow. C) Heatmap showing relative gene expression of glycolysis associated genes from isolated cardiomyocytes of untreated Panx1^fl/fl^ and Panx1^MyHC6^ mice plotted as the Log_2_ (fold change) from Panx1^fl/fl^ mouse. Purple heatmap represents -log(p-value) of two-tailed t-tests comparing fold change of Panx1^fl/fl^ and Panx1^MyHC6^. (Panx1^fl/fl^ N=3 mice, Panx1^MyHC6^ N=3 mice) D) Glucose uptake as measured by 2NBD-glucose fluorescence intensity of control or Panx1-targeted siRNA treated after 1 hour ISO (20μM) stimulation. (N=4) E) Phenogram of bioenergetic profile of control siRNA treated H9c2 cells reveals a shift to more glycolytic phenotype after ISO treatment. siRNA mediated PANX1 knock-down demonstrates a more glycolytic phenotype without ISO stimulation which is only minimally shifted after ISO treatment. F) Quantification of Basal extracellular acidification rate (ECAR) and G) Maximal ECAR of control of Panx1-targeted siRNA treated H9c2 cells after stimulation with either vehicle or ISO (20μM) for 1 hour. (N=5) H) Maximal OCR of control of Panx1-targeted siRNA treated H9c2 cells after stimulation with either vehicle or ISO (20μM) for 1 hour. (N=5) I) Phenogram of ATP production of control siRNA treated H9c2 cells reveals a shift to more glycolytic phenotype after ISO treatment. siRNA mediated PANX1 knock-down demonstrates a more glycolytic phenotype without ISO stimulation which is not shifted after ISO treatment. Data represented as mean ± SEM. Significance determined using One-way ANOVA (D, F-H) with Sidak’s multiple comparison test for post-hoc analysis for comparisons between individual groups. ns = not significant. See also S3 in reference to figure 3.

Pannexin channels have been demonstrated to control cellular metabolism, specifically the regulation of glucose homeostasis and glycolytic metabolism^49–52^. A previous study by Adamson et al. demonstrated that Panx1 channel function is required for insulin-induced glucose metabolism in white adipocytes^53^. To assess whether loss of PANX1 impacted these pathways in cardiomyocytes, we analyzed the genes associated with key glycolytic enzymes in PANX1-deficient cardiomyocytes. Our data demonstrate increased expression of glycolytic pathway genes in cardiomyocytes isolated from Panx1^MyHC6^ mice compared to their floxed controls. Of note, we found a significantly higher levels of the glucose transporter *Slc2a4* (Glut4), which is the predominant isoform in cardiomyocytes (**Figure 3C**). This finding was specific to Glut4 as we do not find any specific changes in Glut1 or Glut3, the other predominate isoforms in cardiomyocytes (S3I).

To determine if the observed transcriptional increase in genes involved in glycolysis impacts cellular metabolism and glucose utilization, we first performed glucose uptake assays to determine if the increase in *Slc2a4* transcripts impacted intracellular glucose content. Intracellular 2NBD-glucose levels were significantly increased after stimulation of H9c2 cells with isoproterenol, which was abrogated after siRNA-mediated knock down of PANX1 (**Figure 3F**).

Next, we performed a glycolytic stress test (GST) using a Seahorse extracellular flux analyzer on H9c2 cells that had been treated with either Panx1 siRNA or control siRNA to determine if increased glucose uptake affected cellular bioenergetics. PANX1 knock-down alone resulted in significantly increased basal and maximal extracellular acidification rate (ECAR). The magnitude of increase was even further increased by treatment with isoproterenol (**Figures 3D, E**, representative trace S3C). Conversely, when we performed a mitochondrial stress test (MST) we found that PANX1 knock-down alone was sufficient to significantly decrease maximal oxygen that PANX1 knock-down drives H9c2 cells to a more glycolytic state, which mimics the shif t seen after isoproterenol treatment in control cells. Moreover, we performed an ATP production assay to determine the contribution of mitochondrial and glycolytic ATP production to the total ATP production rate. We found that isoproterenol treatment of H9c2 rat myoblasts drives an increase in total ATP production, which is predominately glycolytic. PANX1-knock-down similarly significant increases the total ATP production rate through a significantly higher glycolytic ATP production rate compared to control siRNA treated cells (**Figure 3I**, S3F-H). Though cardiomyocyte metabolism and ATP production was significantly changed by PANX1 deletion *in vitro*, intracellular ATP content is not significantly different (S3E). Taken together these data demonstrate a key role for PANX1 channels in regulating glucose uptake and glycolytic rate in H9c2 cardiomyocytes.

### Panx1 deficiency in cardiomyocytes protects against isoproterenol-induced cardiac hypertrophy and cardiac dysfunction in non-ischemic heart failure

To examine the role of Panx1 channels in regulating cardiomyocyte function of the heart under stress, we induced cardiac hypertrophy preceding non-ischemic heart failure using an established protocol where we repeatedly administered isoproterenol via intraperitoneal injection for 14 days at a dose of 15mg/kg/day^41,54^. We established this timepoint as an early measure of disease progression, where cardiac hypertrophy is the predominate phenotype. We performed 2D-echocardiography prior to the beginning of the study to collect baseline measurements and then again after 14 days at the model endpoint (**Figure 4A**). As shown previously, before the start of the treatment Panx1^fl/fl^ and Panx1^MyHC6^ mice displayed no overt differences in cardiac function (Figure 2B-E) or weight (S4A). Repeated isoproterenol administration resulted in significantly increased heart weight in Panx1^fl/fl^ mice, but not in Panx1^MyHC6^ mice (**Figure 4B**, S4B). Cardiomyocyte hypertrophy (as measured by cardiomyocyte cross sectional area by wheat germ agglutinin (WGA) staining) was significantly increased in hearts of in Panx1^fl/fl^ mice compared with Panx1^MyHC6^ mice after isoproterenol treatment (**Figure 4C**).

**Figure 4:**
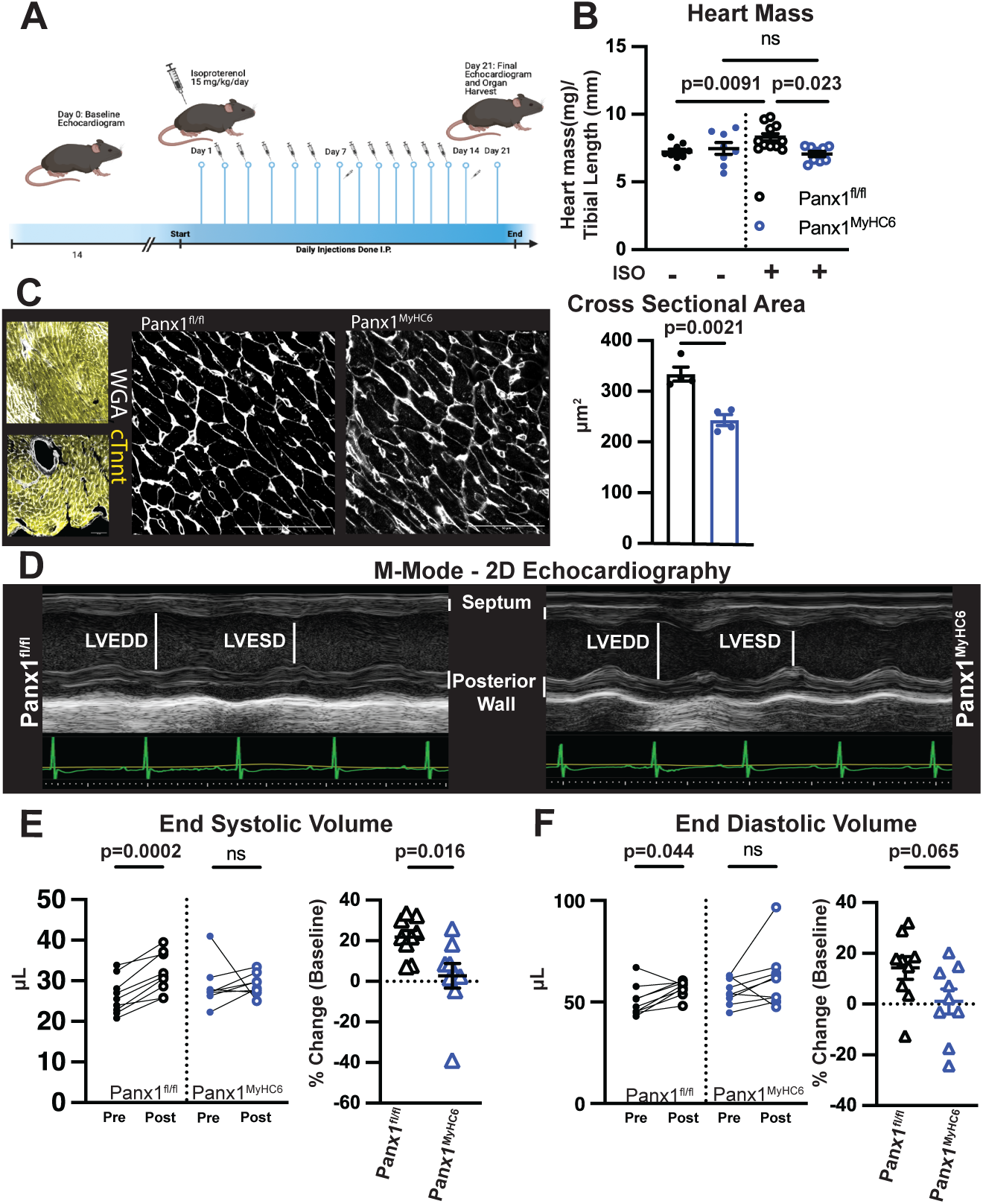
Isoproterenol induced cardiac hypertrophy and dysfunction is abrogated in Panx1^MyHC6^ mice A) Schematic of experimental model of isoproterenol induced non-ischemic heart failure. B) Normalized heart mass from 10–12-week-old Panx1^fl/fl^ or Panx1^MyHC6^ mice treated for 14 days with isoproterenol (15 mg/kg/day), or saline control normalized to tibial length. (Saline: Panx1^fl/fl^ N=9, Panx1^MyHC6^ N= 8; Isoproterenol Panx1^fl/fl^ N=11, Panx1^MyHC6^ N= 9) C) Representative confocal micrographs of wheat germ agglutinin (WGA) staining at 20X (Left) and 60X (Right) from Panx1^fl/fl^ or Panx1^MyHC6^ mice treated with isoproterenol for 14 days. Cardiomyocyte cross sectional area was calculated from WGA. (Panx1^fl/fl^ N=4 mice, Panx1^MyHC6^ N=4, data points represent average of 50 cells per mouse). D) Representative M-mode images of Panx1^fl/fl^ mice or Panx1^MyHC6^ after 14 days of isoproterenol treatment, visualized by 2D echocardiography. E) Left, paired analysis of left ventricle end-systolic volume (μL) pre- and post-isoproterenol treatment; right, percent change from baseline after isoproterenol treatment. (Panx1^fl/fl^ N=9, Panx1^MyHC6^ N=8) F) Left, paired analysis of left ventricle end-diastolic volume (μL) pre- and post-isoproterenol treatment; right, percent change from baseline after isoproterenol treatment. (Panx1^fl/fl^ N=9, Panx1^MyHC6^ N=8) Data represented as mean ± SEM. Significance determined using One-way ANOVA (B with Sidak’s multiple comparison test for post-hoc analysis for comparisons between individual groups; Student’s t-test (C) and Welch’s t-test (E, F) was used for comparison between two groups. Significance of paired data was determined using Mixed effects analysis (E, F) with Sidak’s multiple comparison test for post-hoc analysis for comparisons between timepoints. ns = not significant. See also S4, S5 in reference to figure 4.

Furthermore, treatment with isoproterenol caused significant increases in both end-systolic and end-diastolic volume of the left ventricle in Panx1^fl/fl^ mice as measured by echocardiography (**Figure 4D-F**, S4H). Panx1^MyHC6^ mice did not have significant increases in either metric compared to their baseline and had significant less change compared to Panx1^fl/fl^ age and sex-matched controls (**Figure 4D-F**, S4H). Finally, Panx1^fl/fl^ mice had a trending decrease in ejection fraction after isoproterenol treatment, while Panx1^MyHC6^ had no change (S4G). We determined that this disparity was likely caused by heart-intrinsic functional defects and not systemic vascular or renal damage, as we observed no differences in blood pressure or serum creatinine between Panx1^fl/fl^ and Panx1^MyHC6^ mice after treatment with isoproterenol (S4C-F).

Additionally, to assess cardiac fibrosis found at later stages of non-ischemic heart failure and to examine if Panx1 deficiency would also protect against fibrotic changes and disease progression, we induced non-ischemic heart failure by administration of isoproterenol for 28 days (15/mg/kg/day) by subcutaneous implantation of osmotic pump (S5A). Prolonged isoproterenol treatment caused a significantly greater increase in Collagen 1 (COL1A1) content in the myocardium of Panx1^fl/fl^ mice compared to Panx1^MyHC6^ mice, as determined by immunofluorescence staining (S5B). Moreover, functional analysis by echocardiography revealed that Panx1^MyHC6^ mice had a trending higher percent fractional shortening and a significantly decreased wall thickness compared with age and sex matched Panx1^fl/fl^ controls (S5C,D). These findings indicate a protective effect of cardiomyocyte-specific deletion of PANX1 in the later stages of non-ischemic heart failure.

These data demonstrate that cardiomyocyte-specific deletion of PANX1 is cardioprotective against cardiac hypertrophy and interstitial fibrosis during the progression of non-ischemic heart failure.

### Pathways involved in the immune response and fatty-acid metabolism are down regulated in the hearts of Panx1^MyHC6^ mice during non-ischemic heart failure

To elucidate potential mechanisms that are involved in the protection from cardiac hypertrophy and dysfunction observed in Panx1^MyHC6^ mice, we performed bulk RNA-sequencing on whole hearts of Panx1^fl/fl^ and Panx1^MyHC6^ mice after 14-day administration of isoproterenol. We identified 1592 genes which were significantly differentially expressed between the two genotypes, with 594 genes downregulated and 998 upregulated in Panx1^MyHC6^ mice compared to Panx1^fl/fl^ controls (**Figure 5A**).

**Figure 5:**
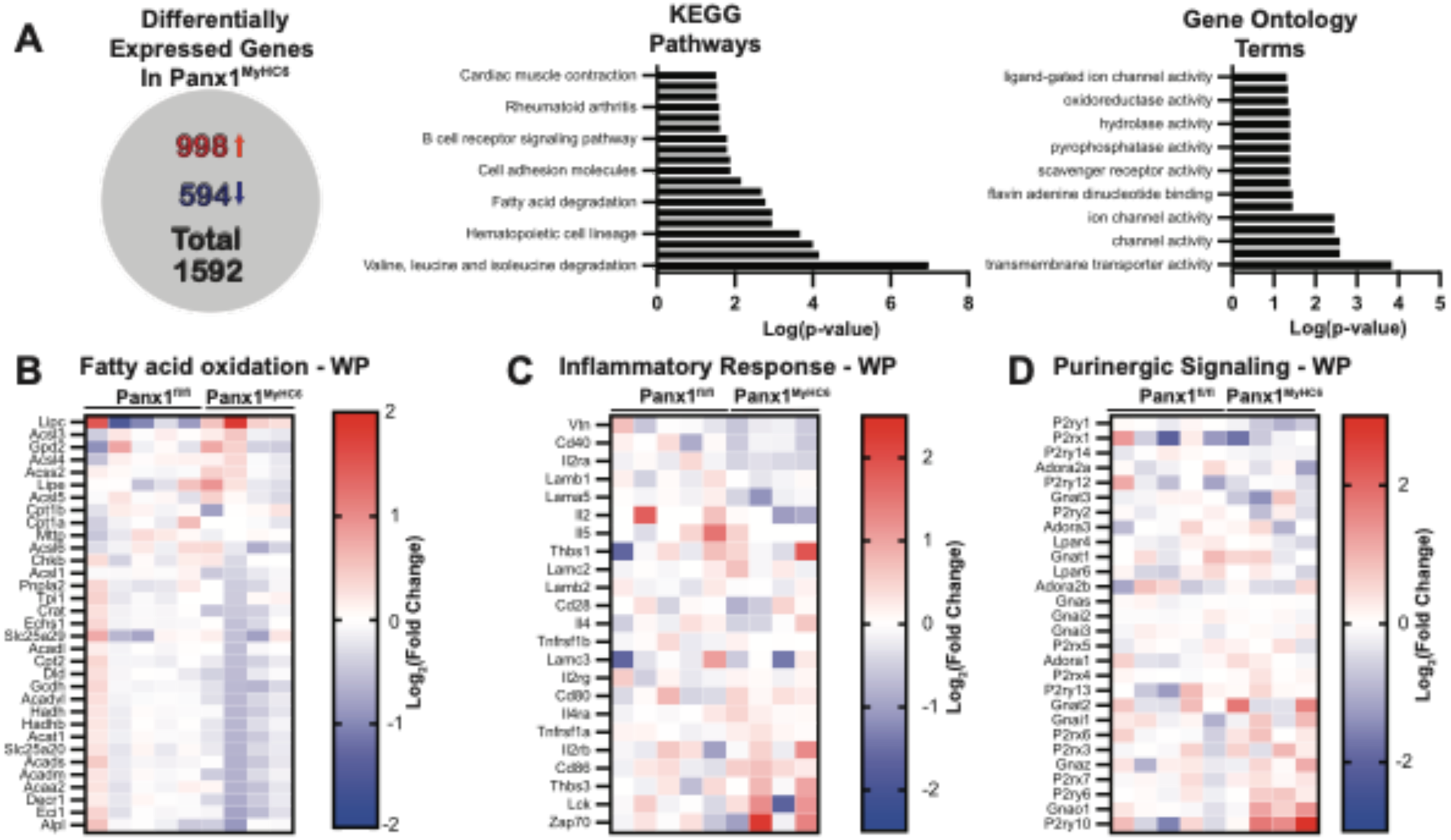
Cardiomyocyte PANX1 knockout leads to transcriptional downregulation of fatty acid metabolism and immune activation pathways during non-ischemic heart failure A) Quantification of differentially expressed genes identified in bulk RNA-Sequencing between whole hearts of Panx1^fl/fl^ and Panx1^MyHC6^ mice after 14 days of isoproterenol treatment. Differentially expressed gene determined by cut off of p < 0.05. Significantly changed, as determined by cut off of p < 0.05, KEGG Pathways and Gene Ontology (GO) Terms identified from differentially expressed genes plotted as ordered by -log(p-value). B) Heatmap demonstrating genes identified in WikiPathways “Inflammatory response”. C) Heatmap demonstrating genes identified in WikiPathways “Fatty acid oxidation and Fatty acid beta-oxidation”. D) Heatmap demonstrating genes identified in WikiPathways “Purinergic signaling”. Gradient plotted as Log_2_(Fold Change) of Panx1^MyHC6^ compared to Panx1^fl/fl^. (Panx1^fl/fl^ N=5, Panx1^MyHC6^ N=4)

KEGG pathway and GO Term analysis on the differentially expressed genes identified regulatory pathway families including “hematopoietic cell lineage”, “cell adhesion molecules”, “B-cell receptor signaling pathway”, and “Rheumatoid arthritis”, all of which suggested significant changes in molecules associated with immune cell recruitment and infiltration. Moreover, we identified “fatty acid degradation”, “fatty acid metabolism”, and “cardiac muscle contraction”, implicating a metabolic shift in the absence of PANX1 channels. Furthermore, identified GO Terms indicated changes in plasma membrane proteins, specifically transporters and ion channels, as well as “oxidoreductase activity”, “scavenger receptor activity”, and “hydrolase activity”, implicating the possible shift in the oxidative stress state.

Further analysis of genes associated with fatty acid beta-oxidation (as identified through WikiPathways) demonstrated overall decreased expression of this group of genes in hearts from Panx1^MyHC6^ mice with non-ischemic heart failure, while some genes such as *Lipc* (encoding the lipase C hepatic type), were expressed at higher levels (**Figure 5B**). Interestingly, a lowered reliance on fatty acid beta-oxidation during ischemic stress has been previously associated with increased cardiomyocyte viability^15^.

Examination of genes involved in the inflammatory response identified lower expression of *Cd40*, *Il2ra*, and *Il2* in hearts of Panx1^MyHC6^ mice compared to their Panx1^fl/fl^ counterparts (**Figure 5C**), suggesting a lower immune cell load in the hearts as CD40 and IL-2 expression are associated with a wide range of leukocyte populations^55^. Additionally, we observed lower expression of *Lamb1* (encoding Laminin subunit beta 1) and *Lama5* (encoding laminin subunit alpha 5) as well as *Vtn* (encoding Vitronectin) suggestive of decreased cell-cell adhesion and fibrosis (**Figure 5C**).

Since PANX1 channel function has been associated with activation of a variety of purinergic receptors^29,56–58^, we examined the expression of purinergic signaling receptors and other associated molecular regulators in our RNA-Seq dataset (WikiPathway “Purinergic signaling”). We found lower expression of *P2yr1*, *P2rx1*, *P2ry14,* and *Adora2a* in hearts of Panx1^MyHC6^ mice (**Figure 5D**) compared to Panx1^fl/fl^ controls. This finding is particularly interesting as *P2rx1, P2yr14*, and *Adora2a* have all been previously associated with chemotaxis and activation of immune cells, particularly neutrophils^59,60^.

Together these data identify that PANX1 deficiency in cardiomyocytes causes significant transcriptomic changes in genes associated with inflammation, immune cell recruitment, metabolism and oxidative phosphorylation, all of which may contribute to the protection of Panx1^MyHC6^ mice from cardiac hypertrophy and progressive heart failure in a model of isoproterenol induced non-ischemic heart failure.

### Neutrophil infiltration is significantly blunted in Panx1^MyHC6^ mice during heart failure induced by isoproterenol

To determine the contribution of cardiomyocytes to the findings of the whole heart, most notably to the inflammatory state in isoproterenol-injured hearts, we performed bulk RNA-Seq on the isolated cardiomyocytes from Panx1^fl/fl^ and Panx1^MyHC6^ mice after 14-days of isoproterenol administration. We identified 1439 differentially expressed genes with the majority being downregulated in Panx1-deficient cardiomyocytes (1037 down, 402 up) (S6A). Further analysis of genes associated with “inflammation response” (WikiPathways) revealed decreased expression of many of the genes identified as being regulatory in inflammation including Zap70, Lack, Fn1, Col1a2, and multiple cytokine receptors (**Figure 6A**). This was corroborated by the finding of the GO Terms “immune response” and “immune system process”, which were significantly enriched with genes downregulated in the isolated cardiomyocyte fraction of Panx1^MyHC6^ mice compared to Panx1^fl/fl^ mice (**Figure 6A**).

**Figure 6:**
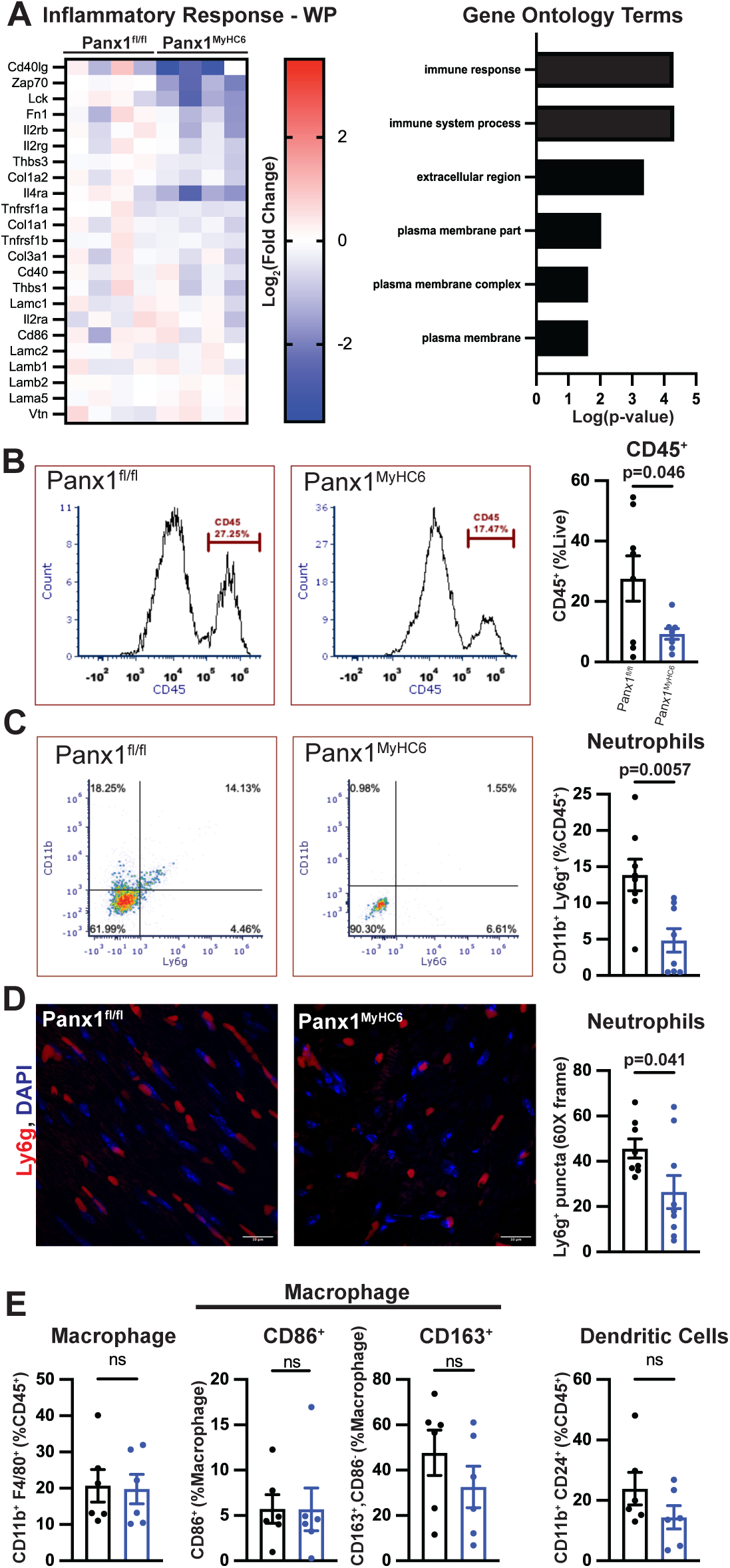
Neutrophil infiltration is significantly blunted in Panx1^MyHC6^ mice during heart failure induced by isoproterenol A) Heatmap demonstrating genes identified in WikiPathway “Inflammatory response”. Gradient plotted as Log_2_(Fold Change) of Panx1^MyHC6^ isolated cardiomyocytes after 14-days of isoproterenol treatment compared to Panx1^fl/fl^. Significantly decreased, as determined by cut off of p < 0.05, GO terms in Panx1^MyHC6^ cardiomyocytes compared with Panx1^fl/fl^ cardiomyocytes. (Panx1^fl/fl^ N=4, Panx1^MyHC6^ N=4 mice). B) Flow cytometric analysis of CD45^+^ cells as a percentage of total live cells isolated from Panx1^fl/fl^ and Panx1^MyHC6^ hearts after 14-days of isoproterenol treatment as identified. Representative histograms of the CD45 populations from live parent gate. (Panx1^fl/fl^ N=8, Panx1^MyHC6^ N=8 mice) C) Flow cytometric analysis of neutrophils (CD11b^+^, Ly6g^+^) as a percentage of the CD45^+^ cells in Panx1^fl/fl^ and Panx1^MyHC6^ hearts after 14-days of isoproterenol treatment. Representative plots of the identified neutrophils from Panx1^fl/fl^ and Panx1^MyHC6^ mice hearts. (Panx1^fl/fl^ N=8, Panx1^MyHC6^ N=8 mice) D) Representative confocal micrographs of Ly6g staining at 60X from Panx1^fl/fl^ and Panx1^MyHC6^ mice treated with isoproterenol for 14 days. Ly6g was used to stain neutrophils; DAPI was used as a nuclear stain. Quantification of number of neutrophils, Ly6g^+^ puncta, per 60X field in whole heart tissue from Panx1^fl/fl^ and Panx1^MyHC6^ mice after 14-days of isoproterenol treatment. (Panx1^fl/fl^ N=8, Panx1^MyHC6^ N=9 mice, 3-5 images per mouse averaged) E) Flow cytometric analysis of macrophages (CD11b^+^, F4/80^+^) as a percentage of CD45^+^ cells from Panx1^fl/fl^ and Panx1^MyHC6^ hearts after 14-days of isoproterenol treatment. Macrophages expressing the co-stimulatory activation marker (CD86^+^) and macrophages

To determine whether the observed inflammatory gene regulation patterns was reflective of perturbed leukocyte recruitment to the hearts of Panx1^MyHC6^ mice, we isolated the non-myocyte fractions from the hearts of Panx1^fl/fl^ and Panx1^MyHC6^ mice after 14-day treatment with isoproterenol and examined leukocyte populations by flow cytometry (representative gating S6B). The number of total leukocytes (CD45^+^) as well as neutrophils (Ly6g^+^, CD11b^+^) were significantly lower in the hearts of Panx1^MyHC6^ mice (**Figure 6B, C,** S6B). These findings were confirmed by immunofluorescence imaging of the left ventricle of hearts from Panx1^fl/fl^ and Panx1^MyHC6^ mice. In accordance with the flow cytometry data, the number of Ly6g^+^ puncta per field in the hearts of Panx1^MyHC6^ mice was significantly lower compared to hearts from Panx1^fl/fl^ mice (**Figure 6D**, S6C).

Of note, there was no significant difference in the total number of macrophages (F4/80^+^, CD11b^+^) present in Panx1^MyHC6^ mouse hearts during non-ischemic heart failure (**Figure 6E**, S6D). Further analysis of macrophage subsets showed no differences in macrophages with the co-stimulatory macrophage activation marker CD86 (F4/80^+^, CD11b^+^, CD86^+^), which was confirmed with qPCR on isolated immune cells from a separate cohort of mice, or macrophages with the anti-inflammatory marker CD163 (F4/80^+^, CD11b^+^, CD163^+^/CD86^-^) (**Figure 6E**, S6D,E). Finally, we observed no differences in dendritic cells (CD11b^+^, CD24^+^) (**Figure 6E**, S6D).

While we did not observe any changes in cell populations, we did find significant decreased in *Il1b and Cxcl2*, M1 macrophage activation markers, expression in isolated immune cells from the myocardium of Panx1^MyHC6^ after 14-day isoproterenol treatment relative to immune cells isolated from Panx1^fl/fl^ mice (S6E). Furthermore, we did not find detectable levels of *Chil3*, an M2 macrophage activation marker, in in isolated immune cells from the myocardium Panx1^fl/fl^, which we were able to observe in isolated immune cells from Panx1^MyHC6^ mice (S6E). Taken together these data, demonstrate that while cell populations of macrophages have not changed the inflammatory and activation state of these populations has significantly shifted.

These data demonstrate a critical role for PANX1 expressed in cardiomyocytes in recruiting immune cells, specifically neutrophils, to the myocardium in response to chronic beta-adrenergic stimulation. This finding is in line with previous work which demonstrates that PANX1-dependent ATP release recruits neutrophils to the endothelium during acute inflammatory disease^61^.

## Discussion

The role of ATP, adenosine, and other purinergic signaling molecules as both autocrine and paracrine signals in the myocardium has long been understood and exploited clinically, using compounds such as clopidogrel and adenosine^62^. However, the underlying mechanisms for release of purinergic compounds in non-ischemic heart failure still need to be investigated^63,64^.

To examine the role of PANX1 in cardiomyocytes, we generated a novel mouse with cardiomyocyte-specific deletion of PANX1 (Panx1^MyHC6^). We found that Panx1^MyHC6^ mice shift the baseline metabolic state of their cardiomyocytes toward a more glycolytic phenotype, concurrent with increased glucose uptake, GLUT4 transporter expression, and glycolytic ATP production. After isoproterenol-induced non-ischemic heart failure, PANX1^MyHC6^ mice are protected from pathological cardiac hypertrophy and dysfunction. We also find that in Panx1^MyHC6^ mouse hearts during isoproterenol-induced cardiac hypertrophy there is a decrease in gene expression of fatty-acid beta oxidation-associated genes. Finally, PANX1 deficiency in cardiomyocytes blunts neutrophil recruitment to the myocardium during isoproterenol-induced non-ischemic heart failure.

Two previous studies by Adamson *et al.* and Senthivinayagam *et al.* found that deletion of PANX1 in either white or brown adipose tissue exacerbated diet-induced obesity, specifically by increasing susceptibility to insulin resistance, establishing that PANX1 plays a critical role in glucose homeostasis in the adipose tissue^28,53^. Since cardiomyocytes are among the most energy-demanding cell types requiring high levels of ATP needed for contraction, we examined the role for PANX1 in control of cardiomyocyte metabolism. We demonstrate that PANX1 deletion induces a metabolic shift in the cardiomyocyte by priming the cardiomyocyte toward a glycolytic state at baseline. Consequently, PANX1-deficient cardiomyocytes are less responsive to stress induced by beta-adrenergic receptor activation. We found a significant increase in *Slc2a4* expression in isolated primary murine cardiomyocytes of Panx1^MyHC6^ mice, which is matched by increased glucose uptake, glycolytic rate, and glycolytic ATP production in PANX1 knock-down H9c2 myoblasts *in vitro*^14^. This glycolytic phenotype has previously been observed in fetal cardiomyocytes and is associated with increased proliferative capacity^11,14^. Furthermore, increased glucose availability and glycolytic rate in unstressed situations leads to better outcomes and is associated with increased regenerative capacity after ischemic/ reperfusion injury^15,65^.

RNA-Sequencing of isolated cardiomyocytes from Panx1^MyHC6^ and Panx1^flfl^ mice with non-ischemic heart failure revealed significantly decreased expression of genes associated with immune system function. Previous work established that PANX1 deletion in endothelial cells decreases the immune cell recruitment to the myocardium after a myocardial infarction, without further specifying a particular immune cell population^35^. We find that deletion of PANX1 in cardiomyocytes decreases overall immune cell recruitment, which was driven specifically by a significant decrease of neutrophils. Immune cell recruitment to the myocardium has been shown to exacerbate cardiomyocyte damage and impairment of myocardial function^7,66^. Therefore, the decrease in immune cells in the myocardium of mice with PANX1-deficient cardiomyocytes suggests that PANX1 may serve to propagate or initiate the sterile immune response in the myocardium during times of chronic cardiac stress^22^. The changes in gene expression reveal an overall decreased chronic inflammatory state of the myocardium in Panx1^MyHC6^ mice after isoproterenol induced heart failure.

While our observations of neutrophil recruitment by PANX1 are in the context of isoproterenol-induced non-ischemic heart failure, PANX1 activation in cardiomyocytes may also be important in ischemic heart failure or after ischemia-reperfusion injury. Future investigations will elucidate if the effects on neutrophil infiltration attributed to cardiomyocyte-specific deletion of PANX1 vary by injury type, which will provide opportunities to develop therapies that are specific to the type of cardiac injury.

ATP is known to serve as recruitment or “find me” signal for immune cells during inflammation^67,68^ and apoptosis^24,25^. While we do not identify the responsible receptors on neutrophils, our RNA sequencing data demonstrate that expression of key purinergic receptors, namely P2X1, P2Y1, and P2Y14, is decreased in the whole heart tissue of Panx1^MyHC6^ mice during isoproterenol-induced heart failure. Of these P2Y1 and P2Y14 can be associated with decreased neutrophil recruitment^39^. Moreover, we cannot rule out other purinergic molecules that could be released through PANX1 potentially driving the neutrophil recruitment. P2Y1 binds ADP which could be generated from local exonucleases acting on the ATP released via PANX1 or could be released from PANX1 itself. Furthermore, we see a decreases in ADORA2A which could be attributed to decrease in neutrophil presence as adenosine released from neutrophils serves to orient neutrophil migration^39^. Additionally, P2Y14 on neutrophils binds UDP-glucose^36^, which can PANX1, including ATP and UDP-glucose, may be necessary for the infiltration of neutrophils into the myocardium during non-ischemic heart failure. ^33,69,70^. However, it is is not completely ^24,35,57^. Historically, treatment of heart failure in patients has focused on therapies which treat the symptoms of the disease, such as edema and dyspnea, rather than directly targeting cardiac function or preventing functional decline of the cardiac muscle. These current clinical therapeutic avenues are limited in impacting the immune cell infiltrate or fibrosis, which are key mechanisms leading to disease progression. Previously clinical trials have identified that improve patient outcomes in both all-cause mortality and sudden cardiac death in patients with established disease over a placebo^33,69,70^. However, the understanding for why this therapy is more beneficial than others in lowering blood pressure is not yet well understood. and decreases cardiac hypertrophy as measured by echocardiography. We find that Panx1^fl/fl^ mice have significant increases in left ventricular end-diastolic and end-systolic volume in isoproterenol induced heart failure. Conversely, Panx1^MyHC6^ do not have any significant increase, and the percent change from baseline in left ventricular volume at both these points is significantly less than their Panx1^fl/fl^ counterparts. With this in mind, we speculate that one immune cell recruitment, as has been shown in multiple mouse models^24,35,57^. Furthermore, modulating cardiomyocyte metabolism to increase glucose uptake may prevent an energy imbalance which is thought to be a driving factor in cardiac muscle function decline. Additionally, the decreased immune cell recruitment may explain why patients with lower natriuretic peptide levels see greater benefit from spironolactone therapy compared to their more diseased counterparts^71^. PANX1 blockade is effective in preventing immune cell infiltration, but has not been shown to reverse previously sustained damage which is more severe in patients with a higher disease burden^7^.

Taken together our study illustrates how PANX1 deletion in cardiomyocytes protects against the progression of non-ischemic heart failure. We demonstrate that PANX1 channel opening can be induced by beta-adrenergic receptor stimulation in cardiomyocytes. Cardiomyocyte-specific deletion in mice (Panx1^MyHC6^) impacts their cardiac glycolytic metabolism, which we attribute to a higher expression of *Slc2a4* and increased glucose uptake. Panx1^MyHC6^ mice are protected from isoproterenol-induced non-ischemic heart failure and cardiac hypertrophy, which we demonstrate to be at least in part due to a decrease in neutrophil recruitment. Our findings identify PANX1 in cardiomyocytes as a therapeutic target to prevent severe cardiac tissue damage in non-ischemic heart failure, which may also be relevant for developing therapies targeted at PANX1 for other forms of heart failure such as HFpEF.

### Limitations

Our study is limited by the ability to discern the metabolite or metabolites released through PANX1 activation *in vivo* which results in neutrophil infiltration. Further, we understand that our model is limited in that it does not examine the role for PANX1 in ischemic injury or the context of ischemia reperfusion injury which can both cause heart failure. We utilize a novel mouse which has constitutive deletion PANX1, therefore we are unable to determine the role for PANX1 in initiation versus the progression of disease. Finally, we use a non-contractile cell line for our *in vitro* studies which does not fully recapitulate the *in vivo* physiology of the myocardium.

## Methods

### Animals

Cardiomyocyte-specific Panx1-deficient mice were generated by crossing *Panx1^loxP/loxP^* (Panx1^fl/fl^) mice^46^ with B6.FVB-Tg(Myh6-cre)2182Mds/J (MyHC6-Cre)^72^ (strain #011038) obtained from the Jackson Laboratory. For breeding purposes, mice hemizygous for the MyHC6 Cre were cross with Panx1^fl/fl^ mice to give littermate mice of Panx1^loxP/loxP^ MyHC6^wt/wt^ (Panx1^fl/fl)^ and Panx1^loxP/loxP^ MyHC6 ^Cre/wt^ (Panx1^MyHC6^). All animals were maintained at the University of Virginia Center for Comparative Medicine in a pathogen-free vivarium facility. Mice were maintained on a 12-hour light/dark cycle and provided ad libitum access to standard rodent chow (Teklad 7912) and water. All experiments performed were approved by the University of Virginia Animal Care and Use Committee (protocol #3444). All experiments were performed using age-matched littermate controls. Genotyping was performed via PCR from tail clip biopsies after proteinase K digestion in DirectPCR tail lysis buffer (Viagen Biotech 102-T). PCR of genomic DNA was performed using Apex Taq Master Mix and allele-specific primers as previously described^73^.

### Isoproterenol induction of heart failure

8–10-week-old male mice were treated daily with isoproterenol or saline vehicle for 14 days to induce heart failure as previously described. Isoproterenol (medchem: HYB0468) was freshly prepared daily in sterile saline prior to injection (15 mg/kg/day) and administered by intraperitoneal injection. Mice were weighed daily to monitor for excessive weight loss and appropriate dose administration.

### Isoproterenol progression of heart failure

For longer-term administration of isoproterenol, osmotic pumps (Alzet 2004) were filled with isoproterenol in sterile saline plus 0.02% acetic acid and mice were weighed prior to implantation to administer 15mg/kg/day according to pump rate specifications. Pumps were equilibrated overnight in sterile saline prior to implantation. 8–10-week-old male mice were anesthetized using isoflurane and implantation sites were exposed after hair removal with Nair hair removal lotion with aloe and lanolin. Mice were administered bupivicane analgesic and sites were sterilized with three washes of betadine and ethanol wipes. Pumps were implanted in a subcutaneous pocket on the back of the mouse and surgical clips were placed for wound closure. Mice were placed on a heating-pad and isoflurane anesthesia was maintained for the length of the procedure. Surgical clips were removed 10-14 days after placement.

### Immunofluorescence staining

Hearts were harvested and fixed for 4 hours in 10% zinc-buffered formalin prior to fixation in 50% ethanol. Tissues were paraffin embedded and cryosectioned to 7μm thickness and mounted. Slides were deparaffinized. Briefly, sections were submerged in Histo-Clear (Electron Microscopy Sciences 64110-01) (3×10 minutes), 100% ethanol (2×3 minutes), 95% ethanol (2×3 minutes), 70% ethanol (1×3 minutes), and PBS (1×5 minutes). Antigen retrieval was performed using citrate-based solution (Vector Laboratories H-330) where slides were submerged in antigen retrieval solution and heated to boiling for 20 minutes, slides were then cooled for 10 minutes at room temperature and 10 minutes at 4°^C^. Tissue sections were then blocked for 1 hour in antibody blocking buffer (0.2% TritonX, 1mL donkey serum) at room temperature. Antibody blocking buffer was removed and replaced with antibody blocking buffer containing primary antibody (1:100) overnight at 4°^C^. Sections were then washed (3xPBS for 5 minutes) and incubated in antibody blocking buffer containing secondary antibody (1:500) for 1 hour at room temperature protected from light. Sections were washed (3xPBS) and counterstained with DAPI (Thermo Fisher Scientific D3571) before mounting. Sections were imaged on an Olympus Fluoview 1000 and are representative images of composite z-stacks. Analysis (thresholding, manual counting, and fluorescence intensity quantification) were performed in ImageJ.

### Quantification of Cardiomyocyte cross-sectional area

Hearts were prepared as indicated in immunofluorescence staining. Sections were stained with cTnnt (1:100 - Abcam ab8295) during primary incubation and DAPI (1:500 - Thermo Fisher D3571) and Wheat-Germ-Agglutinin-Alexa Fluor 647 (5 μg/m – Thermo Fisher W32466) during secondary incubation. Sections were imaged on an Olympus Fluoview 1000 confocal microscope with five images per mouse taken in the left ventricle across two sections. 50 round nucleated cardiomyocytes per mouse were outlined across the images and were analyzed in ImageJ.

### Flow cytometry of infiltrating cardiac leukocytes

Primary infiltrating leukocytes were isolated from cardiac tissue by non-Langendorff perfusion previously described^74^. Briefly, hearts were sequentially perfused with EDTA buffer, perfusion buffer, and digestion buffer containing collagenase until hearts appeared soft. Hearts were then mechanically dissociated with scissors and pipetted up and down. Cells were washed through a 100μm cell strainer to create a single cell suspension. Myocytes were isolated by centrifugation at 120 x g for 5 minutes. Supernatants (containing non-myocyte cells) were transferred to clean tubes and pelleted by centrifugation at 300 x g for 5 minutes. Pellets were treated with ACK lysis buffer for 1 minutes and then resuspended in 1mL of FACS buffer (PBS, 2mM EDTA, and 1%BSA). Splenocytes were isolated by grinding through a 100μm cell strainer and washing with PBS containing 2mM EDTA. Samples were then incubated with ACK lysis buffer for 2 minutes and spun at 400 x g for 5 minutes. All samples were then counted and resuspended at 100μL per 100,000 cells in PBS. Cells were stained with LIVE/DEAD Yellow (1:1000) (Thermo Scientific L34967) for 30 minutes on ice. Fc-receptors were then blocked for 15 minutes using FcBlock (BioRad – BUF041A). Cells were pelleted by centrifugation (300 x g,5 minutes) and resuspended in FACS buffer (100μL per 100,000 cells) and stained with fluorophore-conjugated antibodies (1:100) for 1 hour on ice. See tables for antibodies included below. Cells were washed in PBS (×3) after staining. Flow cytometry collection and deconvolution was performed on an Cytek Northern Lights 3 laser Spectral Flow Cytometer. Automatic deconvolution was performed using single stains generated from splenocytes. Gating and post-hoc analysis was performed with FCS-Express 7.18. Representative gating is presented for all flow cytometry experiments in supplemental figures.

### RNA-Sequencing

Hearts of mice were removed after whole body perfusion with 10mL of PBS, weighed, and snap frozen at -80^C^ prior to analysis. For some experiments, isolated cardiomyocytes were suspended in 1mL of TRIzol^TM^ (Invitrogen) and stored at -80^oC^ for further analysis^74^. After chloroform incubation and centrifugation per manufacture instructions RNA-containing phase was diluted with 70% ethanol 1:1 and RNA isolation was performed using RNeasy Mini Kit (Qiagen). For tissues, Total RNA was isolated using RNeasy Mini Kit (Qiagen) after lysis in RLT buffer with bead homogenization. Total RNA quantity and quality were then assessed on a NanoDrop 2000 spectrophotometer (Thermo Fisher Scientific). Library preparation including quality metrics, sequencing, and clustering were performed by Novogene (Sacramento, CA). Sample integrity was assessed prior to library preparation using the Bioanalyzer 2110 system with an RNA Nano 6000 Assay Kit (Agilent Technologies). One hundred fifty-base paired-end sequencing was performed using an Illumina NovaSeq platform according to manufacture instructions. Fastq files were processed using in-house perl scripts from Novogene to obtain clean reads. Clean reads were aligned to the mouse reference genome using Hisat2 v2.0.5. FeatureCounts was used to calculate read numbers for each gene and fragments of per kilobase of transcript sequence per millions base pairs sequenced (FPKM) were calculated. Differential expression analysis was performed using DESeq2. Differentially expressed genes (DEG) were assigned by criteria of P-value adjusted < 0.05 (-Log(p-value) > 1.3). KEGG pathway analysis and Gene Ontology pathway analysis was performed on DEG using clusterProfiler.

### Echocardiography

8–16-week-old male mice were anesthetized using isoflurane. The chest was exposed after hair removal and a four-lead electrocardiogram (ECG) was collected during the procedure. Images were collected using a VevoView 1100 system. Images were collected in the long-axis (B-mode) and short axis (M-mode) on mice serially throughout the experimental period. Serial collection began prior to disease induction and occurred every two-weeks during disease progression. Images were analyzed using Vevo Lab. Left-ventricle (LV) end-diastolic, end-systolic volume, and wall thickness were assessed, and ejection fraction calculated.

### Blood Pressure Assessments using Radiotelemetry

Blood pressure was measured using telemetry equipment (Data Sciences International, DSI) as previously described^27^. Mice were surgically implanted with radiotelemetry units (PA-C10 or HD-X10). Briefly, while under isoflurane anesthesia, the catheter of a radiotelemetry unit was placed in the left carotid artery and positioned such that the probe reached the aortic arch. The radio-transmitter was placed in a subcutaneous pouch at the right flank. Buprenorphine was used as an analgesic. Mice were allowed to recover for seven days prior to the initiation of recordings. Baseline blood pressure measurements, including systolic pressure, diastolic pressure, mean arterial pressure (MAP) and heart rate, were recorded every minute for a continuous period of 5 days using Dataquest A.R.T. 20 software (DSI). Diurnal (inactive period) MAP was measured during animal’s light cycle (6:00 a.m. to 5:59 p.m.) and nocturnal (active period) MAP was measured during the animal’s dark cycle (6:00 p.m. to 5:59 a.m.).

### Antibodies

Antibodies used in these studies were rabbit monoclonal anti-Panx1 (Cell Signaling - #91137), rabbit monoclonal anti-CCR2 (Abcam – ab273050), rat monoclonal anti-Ly6g (R&D Systems – MAB1037), mouse monoclonal anti-cardiac troponin T (Abcam – ab8295), and wheat germ agglutinin (Invitrogen – W32466). Flow cytometry antibody used include CD45 rat anti- mouse APC eFluor 780 (Invitrogen - 47-0451-80), CD11b rat anti-mouse FITC (Invitrogen - 11- 0112-82), CD163 rat anti-mouse PE (Invitrogen - 12-1631-80), CD86 rat anti-mouse SuperBright 436 (Invitrogen - 62-0869-41), LIVE/DEAD Yellow Cell Stain for 405 (L34967), Ly6g rat anti-mouse PE-Cyanine5 (Invitrogen - 15-9668-80), F4/80 anti-mouse APC (eBioscience 17-4801-82), CD64 anti mouse PerCP-710 (Invitrogen – 46-0641-80), CD24 anti mouse PE-Cyanine7 (Invitrogen 25- 0242-82).

### Cell culture

H9c2 rat myoblasts were obtained from ATTC and maintained in high-glucose DMEM supplemented with 10% heat-inactivated FBS, 15 mM sodium bicarbonate (Glibco), 1mM sodium pyruvate (Glibco), and 1% Pen-Strep. Cells were passaged when they reached 70% confluency on tissue culture (TC) treated dishes using 0.25% trypsin (Glibco).

### siRNA-Mediated Knock-Down of Panx1

H9c2 cells in 12 or 24 well TC treated plates or 10cm TC treated dishes were washed in sterile PBS and then incubated with OptiMEM (Glibco). Cells were treated with RNAiMax Lipofectamine and siRNA against Panx1 or negative control silencer #1 (Ambion - 197003) per the manufacturer’s instructions. Experiments were performed 72 hours after transfection with the siRNA.

### RNA isolation and quantitative reverse transcription PCR

RNA was isolated from approximately 75,000 H9c2 rat myoblasts or isolated cardiomyocytes from one mouse heart and lysed in RLT lysis buffer. RNA was isolated using the RNeasy Mini Kit (QIAGEN) according to the manufacturer’s instructions. RNA was quantified using a NanoDrop spectrophotometer (Thermo Fisher Scientific), and complementary DNA (cDNA) libraries were generated using the iScript cDNA Synthesis Kit (Bio-Rad). Quantitative reverse transcription PCR was performed using TaqManFast Advanced Master Mix (Thermo Fisher Scientific) and TaqMan primers (mouse *Panx1* mm0045091, mouse *Panx2* mm01308054, mouse *Pan×3* mm00552586, mouse *Gja1* mm00439105, mouse *Gja5* mm00433619– Invitrogen) (Rat *Panx1* Rn01447976_m1 – Thermo Fisher Scientific) for genes of interest on a CFX Connect real-time PCR instrument (Bio-Rad). Relative gene expression was determined using *18S* (4333760 – Thermo Fisher Scientific) or *Hprt* (mm01545399 – Invitrogen) as a housekeeping gene. Quantitative reverse transcription PCR was performed using SensiMix SYBR Green reagent and primer pairs for genes of interest on a CFV Connect real-time PCR instrument where TaqMan primers were not used. Relative gene expression was determined using *Hprt* as a housekeeping gene. Primer sequences were verified against genes of interest using NCBI Primer-BLAST.

### ATP release measurements

H9c2 cells were plated at 80,000 cells per well in a 48 well dish and put in serum free media overnight prior to experimental treatment. Cells were incubated with Panx1 inhibitors (spironolactone 50 μM in serum free media) or vehicle control for 1 hour prior to stimulation. ARL (Tocris 1283) (300 μM) was added for 30 minutes prior to stimulation. Cells were then stimulated with vehicle (PBS), isoproterenol (2 or 20 μM in sterile PBS), or phenylephrine positive control (10 μM) for 15 minutes. One-third of the media in the well was carefully collected with care taken to not bump cells in the bottom of the well and placed on ice. ATP concentrations were assessed by CellTiterGlo (Promega G7571) and compared to standard curve prepared by combining serum free H9c2 media with ATP disodium salt (Tocris 3245).

### Dye Uptake Assay

H9c2 cells were plated at 1000,000 cells per well in a 12 well dish and incubated with Panx1 inhibitors (spironolactone 50 μM, carbenoxolone 50 μM, or probenecid 1mM) or vehicle control for 1 hour prior to stimulation. Cells were stimulated with vehicle (PBS), isoproterenol (2 or 20 μM in sterile PBS), or phenylephrine positive control (10 μM) for 1 hour at the same time as Yo-Pro-1 dye (Thermo Fisher Scientific Y3603) (1μM) was added to the media. Media was removed and cells were washed prior to detachment with trypsin (0.25%). Cells were spun at 400 x g for 5 minutes and resuspended in PBS with LIVE/DEAD Yellow (1:1000) for 30 minutes. Cells were pelleted and washed in Annexin V binding buffer three times and then resuspended in Annexin V AlexaFluor 568 (Thermo Fisher Scientific A13202) (1:500) (AV) for 10 minutes. Cells were washed and resuspended in FACS buffer as described above. Cells were analyzed by flow cytometry on an Attune NxT. Single stain controls were performed using H9c2 cells for initial intensity. FMO were collected and used for deconvolution prior to gating. Cells were gated for LIVE/DEAD Yellow and AV negative and the percentage of cells positive for Yo-Pro-1 was assessed as compared to vehicle stimulation.

### Glycolytic Stress Test,Mitochondrial Stress Test, and ATP Production assay

After treatment with control or Panx1-targeting siRNA as described above, H9c2 rat myoblasts were plated (40,000 cells per well) in complete medium in XFe96 cell culture microplate and allowed to settle for one hour at room temperature before overnight culture. The following day cells were treated with ISO (20μM) for 1 hour in complete medium. Immediately prior to the assay the media was changed to glycolytic stress test medium (Sigma Aldrich D5030) supplemented with 143 mM NaCl and 2mM L-glutamine (Gibco 25030 – 081), mitochondrial stress test medium (Corning 50-003-PB), or ATP production medium (Agilent 103575-100). Glycolytic activity was assessed by measurement of extracellular acidification rate on a Seahorse XFe96 instrument (Agilent Technologies). The rate of pH change was measured every 13 minutes for a 3-minute interval before sequential challenge with 1) 20mM glucose (Sigma Aldrich D9434), 2) 1 μM oligomycin (Sigma Aldrich 75351), and 3) 80 mM 2-deoxyglucose (Sigma Aldrich D8375). Basal ECAR was measured as post-glucose ECAR minus post 2-DG ECAR, and maximal ECAR was measured as post-oligomycin ECAR minus post-2DG ECAR. Mitochondrial activity was assessed by measurement of O_2_ consumption rate on a Seahorse XFe96 instrument (Agilent Technologies). The rate of O_2_ change was measured every 13 minutes for a 3-minute interval before sequential challenge with 1) 1 μM oligomycin (Sigma Aldrich 75351), 2) 2 μM BAM15 (Cayman Chemical Company 17811) and 3) 10 μM antimycin A (Sigma-Aldrich, A8674) and 10 μM rotenone (Sigma-Aldrich, R88751G). Maximal OCR was measured as post-BAM15 OCR minus post-antimycin A/rotenone OCR. ATP production was assessed by measurement of proton extrusion rate (PER) on a Seahorse XFe96 instrument (Agilent Technologies). The PER was measured every 13 minutes for a 3-minute interval before sequential challenge with 1) oligomycin and 2) antimycin A and rotenone according to manufacture instructions.

### 2-NDB Glucose Uptake

H9c2 rat myoblasts were plated in a black, clear bottom 96 well plate and allowed to adhere overnight in complete media. Cells had been cultured with either control or Panx1 siRNA for 48 hours in OptiMem media prior to plating. The following day cells were treated with ISO (20μM) for 1 hour or vehicle control. After stimulation media was removed and replaced with glucose free DMEM as described above in the presence of 2-NDB glucose for 4 hours. The uptake of fluorescent 2-NBDG was assessed according to manufacturer’s instructions (Cayman Chemicals - Glucose Uptake Cell-Based Assay Kit) on a Synergy HTX plate reader (BioTek).

### CytoTox Cell Death Assay

H9c2 rat myoblasts were plated in a clear bottom 96 well plate and allowed to adhere overnight in complete phenol red free media. Cells had been cultured with either control or Panx1 siRNA for 48 hours in OptiMem media prior to plating. The following day cells were treated with ISO (2,20μM) or phenylephrine (10 μM) for 15 minutes or vehicle control. After stimulation media was removed and LDH presence in the media was assessed according to manufacturer’s instructions (Promega-CytoTox Non-radioactive cytotoxicity assay) on a Synergy HTX plate reader (BioTek).

### Serum Creatinine

Whole blood was collected from mice via cardiac puncture and placed into serum separating tubes (BD Microtainer – 365967) for 30 minutes at room temperature. Serum was separated by centrifugation (1500 x g, 5 minutes) and snap frozen on dry ice. Serum creatinine was measured at 520 nm after addition of sodium picrate according to manufacturer’s instructions (Pointe Scientific -Creatinine Reagent Set C75391250) using a known standard (Pointe Scientific C75391250).

### Statistics

Statistical analyses were performed in Prism 9 (GraphPad). The presence of statistical outliers was assessed by a ROUT test (2%), and identified outliers were excluded from analysis. Statistical outliers were identified for data show in Figure 4E, F, which were excluded from analysis. Statistical tests were performed with two-tailed analysis. Comparisons between two groups was conducted by Welch’s or Student’s t-test, while comparisons between more than two groups were made by one-way analysis of variance or mixed-effects analysis. Sidak’s multiple comparisons test was performed as post-hoc analysis where appropriate. Mixed effects analysis was used for paired data with more than two groups. A p-value of less than 0.05 was considered statistically significant. Data are represented as mean ± the standard error of the mean. Image analysis was performed on 3-5 averaged images per mouse across a minimum of two cardiac sections. Data points shown for imaging are representative of the average across a mouse, with each data point representative of individual mice.

### Data Availability

Sequencing data reported in this manuscript are deposited in the NCBI Gene expression Omnibus. All data needed to evaluate the conclusions of this manuscript are present in the manuscript, present in the supplementary figures, or available upon request to the corresponding author. Graphics were prepared using BioRender. Panx1^MyHC6^ mice can be provided by N.L. upon completion of a materials transfer agreement with the University of Virginia School of Medicine. Requests may be submitted to N.L.

## Acknowledgements

We thank the members of the Pannexin Interest Group for their input and expertise throughout the project. We thank the Cardiovascular Research Center histology core, and the Flow Cytometry Core Facility at the University of Virginia School of Medicine for their expertise and resources.

