## Supplemental figures and legends for "Pannexin 1 Channels Control Cardiomyocyte Metabolism and Neutrophil Recruitment During Non-Ischemic Heart Failure"

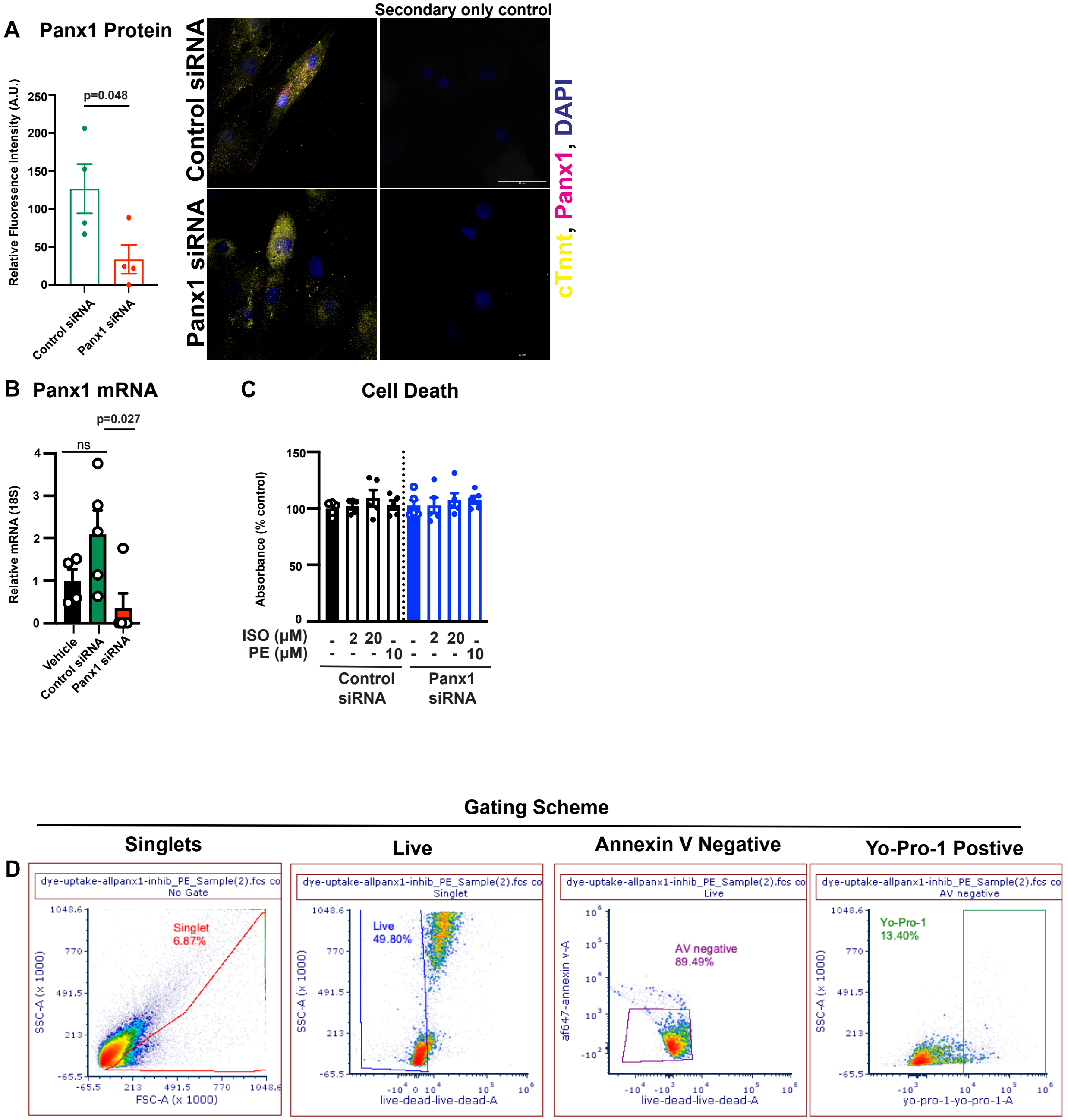


Supplementary Figure 1:

A) Left, quantification from confocal micrographs of Panx1 in control siRNA and Panx1-targeted siRNA in H9c2 cells after 72 hours of treatment for mean fluorescence intensity. Cardiac troponin T (cTnnt) was used to stain cardiomyocytes; DAPI was used as a nuclear stain. Right, secondary only controls included for cTnnt and Panx1 staining. (N=4)

B) Relative mRNA expression of *Panx1* (18S) in vehicle, control siRNA, and Panx1-targeted siRNA treated H9c2 cells after 48 hours of treatment (Vehicle N=4, Control siRNA N=5, Panx1 siRNA N=5).

C) H9c2 rat myoblasts were transfected with control or Panx1-targeted siRNA for 72 hours. LDH was measured after stimulation with ISO (2 or 20 μM), phenylephrine (10 μM), or vehicle control as a proxy for cell death. All data was normalized to the absorbance of the vehicle control. (N=5)

D) Gating scheme for the identification of Yo-Pro-1 positive H9c2 rat myoblasts.

Data represented as mean $\pm$ SEM. Significance determined using Student's t-test (A) or One-way ANOVA (B) with Sidak's multiple comparison test for post-hoc analysis for comparisons between individual groups. * $p < 0.05$, ns = not significant

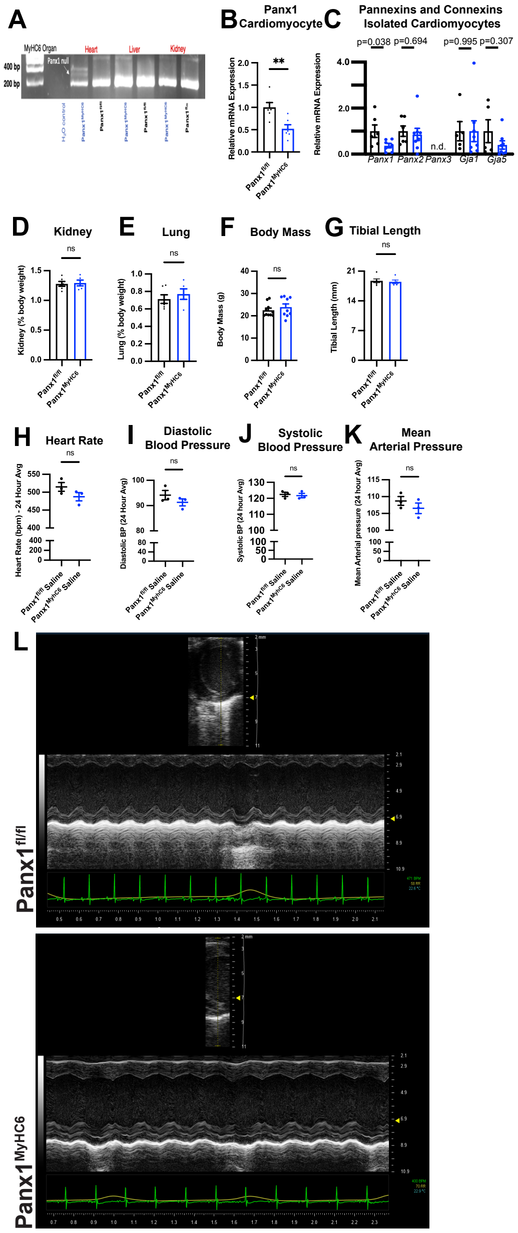


Supplementary Figure 2:

A) Genomic DNA from PCR for Panx1 in heart of Panx1$^{fl/fl}$ and Panx1$^{MyHC6}$ mouse with null band present in Panx1$^{MyHC6}$ heart. Panx1 null band absent from genomic DNA from PCR for Panx1 in heart of Panx1$^{fl/fl}$ and from liver or kidney of Panx1$^{fl/fl}$ and Panx1$^{MyHC6}$ mice.

B) Relative mRNA expression of *Panx1* (Hprt) in isolated cardiomyocytes from untreated Panx1$^{fl/fl}$ and Panx1$^{MyHC6}$ mice. (Panx1$^{fl/fl}$ N=6, Panx1$^{MyHC6}$ N=6)

C) Relative mRNA expression of *Panx1, Panx2, Panx3, Gja1* (Cx43)*, Gja5* (Cx40) in isolated cardiomyocytes from untreated Panx1$^{fl/fl}$ and Panx1$^{MyHC6}$ mice. (Panx1, Panx2: Panx1$^{fl/fl}$ N=6, Panx1$^{MyHC6}$ N=8; Gja1, Gja5: Panx1$^{fl/fl}$ N=5, Panx1$^{MyHC6}$ N=8)

D) Kidney weight as a percentage of body weight of 12-week-old male untreated Panx1$^{fl/fl}$ and Panx1$^{MyHC6}$ mice. (Panx1$^{fl/fl}$ N=6, Panx1$^{MyHC6}$ N=5)

E) Lung weight as a percentage of body weight of 12-week-old male untreated Panx1$^{fl/fl}$ and Panx1$^{MyHC6}$ mice. (Panx1$^{fl/fl}$ N=6, Panx1$^{MyHC6}$ N=5)

F) Body mass of 12-week-old male untreated Panx1$^{fl/fl}$ and Panx1$^{MyHC6}$ mice. (Panx1$^{fl/fl}$ N=10, Panx1$^{MyHC6}$ N=9)

G) Tibial length of 12-week-old male untreated Panx1$^{fl/fl}$ and Panx1$^{MyHC6}$ mice measured with manual calipers. (Panx1$^{fl/fl}$ N=6, Panx1$^{MyHC6}$ N=5)

H) 24-hour average heart rate measured by radio telemetry for 5 days averaged in 12-week-old male saline treated (14 days i.p.) Panx1$^{fl/fl}$ and Panx1$^{MyHC6}$ mice. (Panx1$^{fl/fl}$ N=3, Panx1$^{MyHC6}$ N=3)

I) 24-hour average diastolic blood pressure measured by radio telemetry for 5 days averaged in 12-week-old male saline treated (14 days i.p.) Panx1$^{fl/fl}$ and Panx1$^{MyHC6}$ mice. (Panx1$^{fl/fl}$ N=3, Panx1$^{MyHC6}$ N=3)

J) 24-hour average systolic blood pressure measured by radio telemetry for 5 days averaged in 12-week-old male saline treated (14 days i.p.) Panx1$^{fl/fl}$ and Panx1$^{MyHC6}$ mice. (Panx1$^{fl/fl}$ N=3, Panx1$^{MyHC6}$ N=3)

K) 24-hour average mean arterial pressure measured by radio telemetry for 5 days averaged in 12-week-old male saline treated (14 days i.p.) Panx1$^{fl/fl}$ and Panx1$^{MyHC6}$ mice. (Panx1$^{fl/fl}$ N=3, Panx1$^{MyHC6}$ N=3)

L) Representative images of short-axis B-mode and M-mode 2D echocardiography of Panx1$^{fl/fl}$ and Panx1$^{MyHC6}$ mice prior to the beginning of isoproterenol injections.

Data represented as mean $\pm$ SEM. Significance determined using Student's t-test (B-J).

ns = not significant

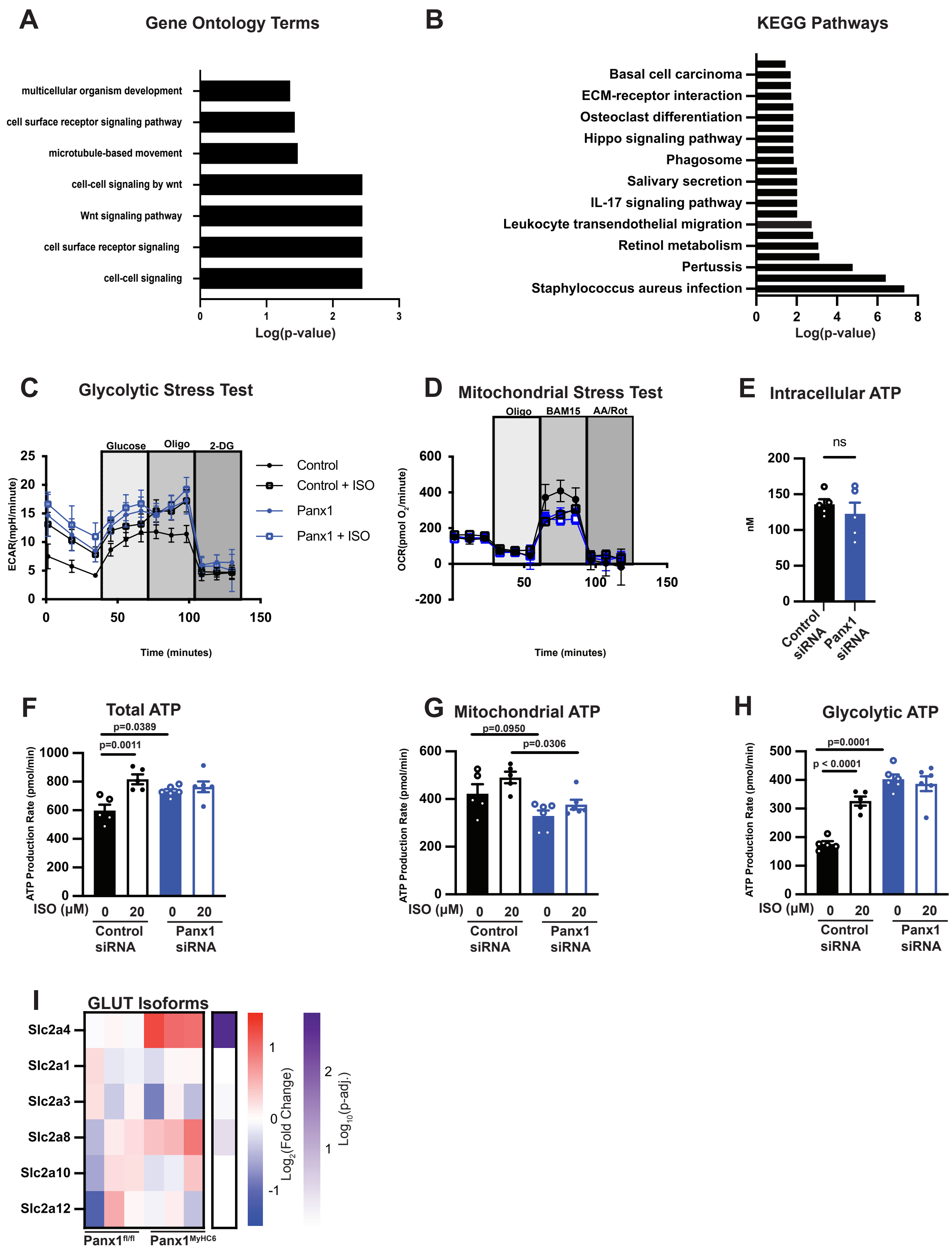
A
Gene Ontology Terms
multicellular organism development
cell surface receptor signaling pathway
microtubule-based movement
cell-cell signaling by wnt
Wnt signaling pathway
cell surface receptor signaling
cell-cell signaling
Log(p-value)
B
KEGG Pathways
Basal cell carcinoma
ECM-receptor interaction
Osteoclast differentiation
Hippo signaling pathway
Phagosome
Salivary secretion
IL-17 signaling pathway
Leukocyte transendothelial migration
Retinol metabolism
Pertussis
Staphylococcus aureus infection
Log(p-value)
C
Glycolytic Stress Test
Glucose
Oligo
2-DG
ECAR(mpH/minute)
Control
Control + ISO
Panx1
Panx1 + ISO
Time (minutes)
D
Mitochondrial Stress Test
Oligo
BAM15
AA/Rot
OCR(pmol O2/minute)
Time (minutes)
E
Intracellular ATP
ns
nM
Control siRNA
Panx1 siRNA
F
Total ATP
p=0.0389
p=0.0011
ATP Production Rate (pmol/min)
ISO (μM)
Control siRNA
Panx1 siRNA
G
Mitochondrial ATP
p=0.0950
p=0.0306
ATP Production Rate (pmol/min)
ISO (μM)
Control siRNA
Panx1 siRNA
H
Glycolytic ATP
p=0.0001
p < 0.0001
ATP Production Rate (pmol/min)
ISO (μM)
Control siRNA
Panx1 siRNA
I
GLUT Isoforms
Slc2a4
Slc2a1
Slc2a3
Slc2a8
Slc2a10
Slc2a12
Panx1fl/fl
Panx1MyHC6
Log2(Fold Change)
Log10(p-adj.)

Revised Supplementary Figure 3:

A) Significant, as determined by a cutoff of $p < 0.05$, GO Terms identified from the differential gene expression in isolated cardiomyocytes of untreated Panx1$^{fl/fl}$ and Panx1$^{MyHC6}$ mice. (Panx1$^{fl/fl}$ N=3, Panx1$^{MyHC6}$ N=3)

B) Significant, as determined by a cut off of $p < 0.05$, KEGG pathways identified from the differential gene expression in isolated cardiomyocytes of untreated Panx1$^{fl/fl}$ and Panx1$^{MyHC6}$ mice. (Panx1$^{fl/fl}$ N=3, Panx1$^{MyHC6}$ N=3)

C) H9c2 cells were transfected with either control or Panx1-targeting siRNA and treated with either vehicle or ISO (20µM) for 1 hour. Extracellular acidification was measured after the addition of glucose, oligomycin (oligo), and 2-deoxyglucose (2-DG). (N=5)

D) H9c2 cells were transfected with either control or Panx1-targeting siRNA and treated with either vehicle or ISO (20µM) for 1 hour. Oxygen consumption rate was measured after the addition of oligomycin (oligo), BAM15, and antimycin A and rotenone. (N=3-6)

E) H9c2 cells were transfected with either control or Panx1-targeting siRNA for 72 hours and intracellular concentration of ATP was measured. (N=5)

H9c2 cells were transfected with either control or Panx1-targeting siRNA and treated with either vehicle or ISO (20µM) for 1 hour. F) Total ATP, G) Mitochondrial ATP, H) Glycolytic ATP production was measured by proton extrusion rate after the addition of oligomycin (oligo), and antimycin A and rotenone. (N=5-6)

I) Heatmap showing relative gene expression of predominate GLUT isoforms in isolated cardiomyocytes of untreated Panx1$^{fl/fl}$ and Panx1$^{MyHC6}$ mice plotted as the $Log_2$ (fold change) from Panx1$^{fl/fl}$ mouse. Purple heatmap represents -log(p-value) of two-tailed ttests comparing fold change of $Panx1^{fl/fl}$ and $Panx1^{MyHC6}$. ($Panx1^{fl/fl}$ N=3 mice, $Panx1^{MyHC6}$ N=3 mice)

Data represented as mean ± SEM. Significance determined using Student's t-test (E). One-way ANOVA (F-H) with Sidak's multiple comparison test for post-hoc analysis for comparisons between individual groups. ns = not significant

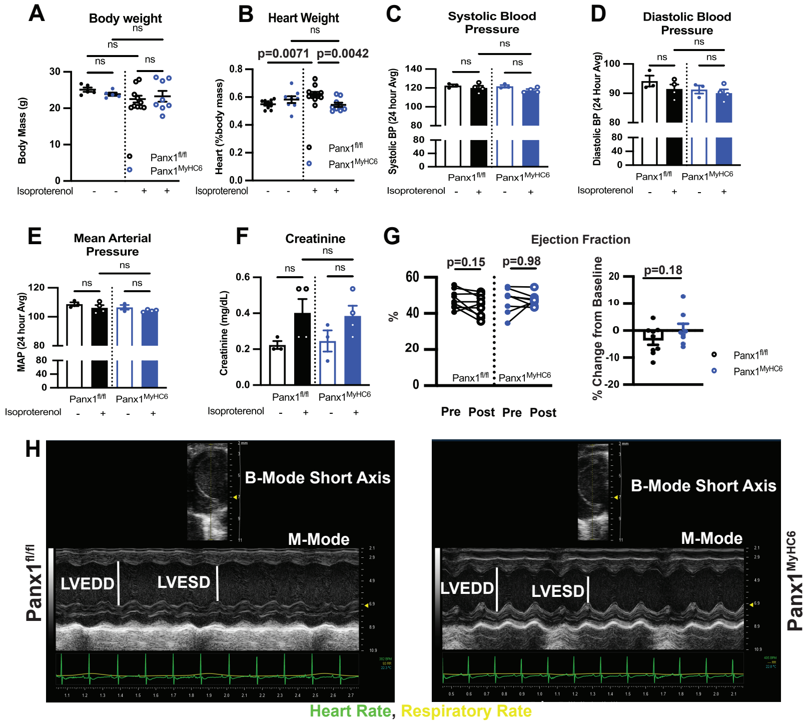


Revised Supplementary Figure 4:

A) Body weight of Panx1$^{fl/fl}$ and Panx1$^{MyHC6}$ mice treated with isoproterenol (15 mg/kg/day) or saline control for 14 days. (Saline: Panx1$^{fl/fl}$ N=9, Panx1$^{MyHC6}$ N= 8; Isoproterenol Panx1$^{fl/fl}$ N=11, Panx1$^{MyHC6}$ N= 9)

B) Normalized heart weight as a percentage of body weight of Panx1$^{fl/fl}$ and Panx1$^{MyHC6}$ mice treated with isoproterenol (15 mg/kg/day) or saline control for 14 days. (Saline: Panx1$^{fl/fl}$ N=9, Panx1$^{MyHC6}$ N= 8; Isoproterenol Panx1$^{fl/fl}$ N=11, Panx1$^{MyHC6}$ N= 9).

C) 24-hour average systolic blood pressure measured by radio telemetry for 5 days averaged in Panx1$^{fl/fl}$ and Panx1$^{MyHC6}$ mice treated with isoproterenol (15 mg/kg/day) or saline control for 14 days. (Saline: Panx1$^{fl/fl}$ N=3, Panx1$^{MyHC6}$ N= 3; Isoproterenol Panx1$^{fl/fl}$ N=4, Panx1$^{MyHC6}$ N= 4)

D) 24-hour average diastolic blood pressure measured by radio telemetry for 5 days averaged in Panx1$^{fl/fl}$ and Panx1$^{MyHC6}$ mice treated with isoproterenol (15 mg/kg/day) or saline control for 14 days. (Saline: Panx1$^{fl/fl}$ N=3, Panx1$^{MyHC6}$ N= 3; Isoproterenol Panx1$^{fl/fl}$ N=4, Panx1$^{MyHC6}$ N= 4)

E) 24-hour average mean arterial pressure measured by radio telemetry for 5 days averaged in Panx1$^{fl/fl}$ and Panx1$^{MyHC6}$ mice treated with isoproterenol (15 mg/kg/day) or saline control for 14 days. (Saline: Panx1$^{fl/fl}$ N=3, Panx1$^{MyHC6}$ N= 3; Isoproterenol Panx1$^{fl/fl}$ N=4, Panx1$^{MyHC6}$ N= 4)

F) Serum creatinine in Panx1$^{fl/fl}$ and Panx1$^{MyHC6}$ mice treated with isoproterenol (15 mg/kg/day) or saline control for 14 days. (Saline: Panx1$^{fl/fl}$ N=3, Panx1$^{MyHC6}$ N= 3; Isoproterenol Panx1$^{fl/fl}$ N=4, Panx1$^{MyHC6}$ N= 4)

G) Left, paired analysis of left ventricle ejection fraction (%) pre- and post-isoproterenol treatment; right, percent change from baseline after isoproterenol treatment. (Panx1$^{fl/fl}$ N=9, Panx1$^{MyHC6}$ N=8)

H) Representative images of short-axis B-mode and M-mode 2D echocardiography of Panx1$^{fl/fl}$ and Panx1$^{MyHC6}$ mice treated with isoproterenol (15 mg/kg/day) for 14 days. Data represented as mean $\pm$ SEM. Significance determined using One-way ANOVA (A-F) with Sidak’s multiple comparison test for post-hoc analysis for comparisons between individual groups. Significance of paired data was determined using Mixed effects analysis (G) with Sidak’s multiple comparison test for post-hoc analysis for comparisons between timepoints. ns = not significant

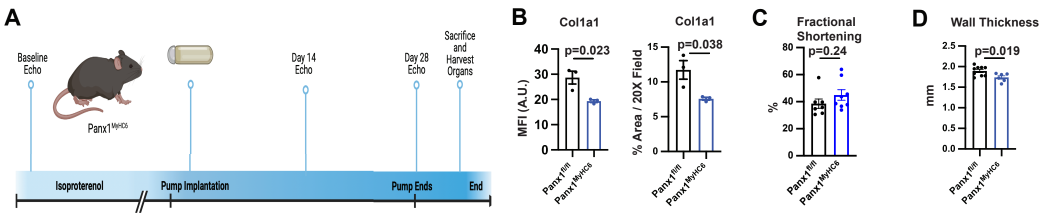


Revised Supplementary Figure 5:

A) Schematic of experimental design for osmotic pump implantation for isoproterenol treatment for 28 days continuous administration (15 mg/kg/day) in Panx1$^{fl/fl}$ and Panx1$^{MyHC6}$ mice.

B) Quantification of confocal micrographs of Collagen 1a1 for intensity (mean fluorescence intensity) and percent area (20X field) after 28 days of isoproterenol administration in 8–10-week-old male Panx1$^{fl/fl}$ and Panx1$^{MyHC6}$ mice. (Panx1$^{fl/fl}$ N=3, Panx1$^{MyHC6}$ N=3)

C) Fractional shortening as measured by echocardiography after 28 days of isoproterenol administration in 8–10-week-old male Panx1$^{fl/fl}$ and Panx1$^{MyHC6}$ mice. (Panx1$^{fl/fl}$ N=7, Panx1$^{MyHC6}$ N=8)

D) Wall thickness of combined anterior and posterior wall of the left ventricle (mm) after 28 days of isoproterenol administration in 8–10-week-old male Panx1$^{fl/fl}$ and Panx1$^{MyHC6}$ mice. (Panx1$^{fl/fl}$ N=7, Panx1$^{MyHC6}$ N=8)

Data represented as mean $\pm$ SEM. Significance determined using Student's t-test (B-D).

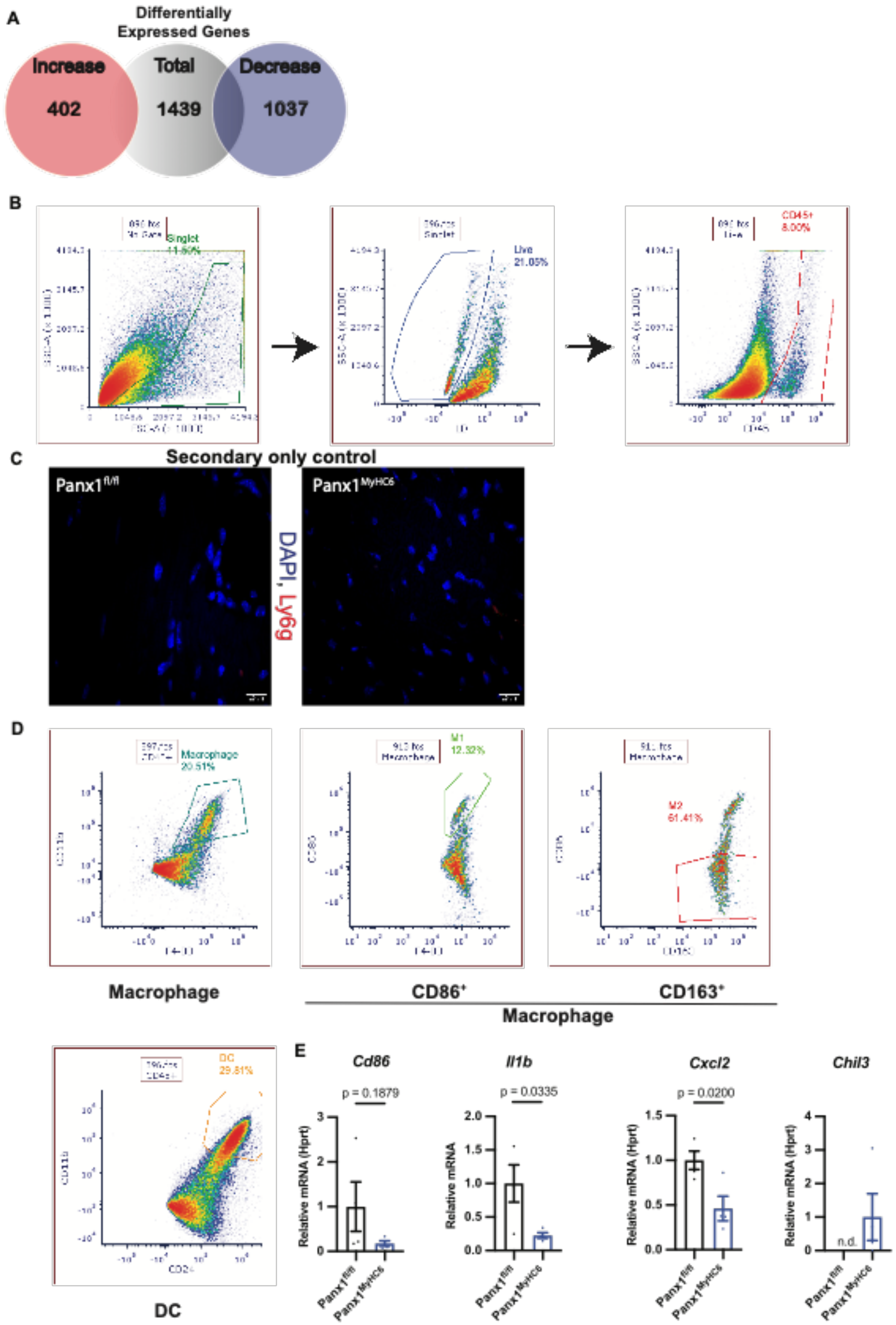
A
Differentially Expressed Genes
Increase
402
Total
1439
Decrease
1037
B
Singlet
Live
21.05%
CD45+
8.00%
C
Secondary only control
Panx1 fl/fl
Panx1 MyHC6
DAPI, Ly6g
D
Macrophage
20.51%
M1
12.32%
M2
61.41%
Macrophage
CD86+
CD163+
Macrophage
DC
29.81%
DC
E
Cd86
p = 0.1879
Il1b
p = 0.0335
Cxcl2
p = 0.0200
Chil3
n.d.
Relative mRNA (Hprt)
Relative mRNA
Panx1 fl/fl
Panx1 MyHC6

Revised Supplemental Figure 6:

A) Venn Diagram of the quantification of significantly increased (left, red), total (middle), and decreased (right, blue) differentially expressed genes, as determined by a cut off of $p < 0.05$, in isolated cardiomyocytes from $Panx1^{fl/fl}$ and $Panx1^{MyHC6}$ mice treated with isoproterenol for 14 days (15 mg/kg/day).

B) Gating scheme and representative gates for the identification of $CD45^{+}$ cells from the non-myocyte fraction of mouse myocardial tissue.

C) Confocal micrographs of secondary only controls from cardiac sections from $Panx1^{fl/fl}$ and $Panx1^{MyHC6}$ mice treated with isoproterenol (15mg/kg/day) for 14 days. Sections were stained with secondary antibody Goat anti-donkey Alexa Fluor 647; DAPI was used as a nuclear stain.

D) Flow cytometric analysis with representative gates for the identification of macrophages ($CD11b^{+}$, $F4/80^{+}$), macrophages expressing the co-stimulatory activation marker CD86 ($CD86^{+}$), macrophages expressing the anti-inflammatory M2 marker ($CD163^{+}$). Flow cytometric analysis with representative gates for the identification of dendritic cells ($CD11b^{+}$, $CD24^{+}$) from the $CD45^{+}$ cells of the non-myocyte fraction of the myocardium of $Panx1^{fl/fl}$ and $Panx1^{MyHC6}$ mice treated with isoproterenol for 14 days (15 mg/kg/day).

E) Relative mRNA expression (Hprt) of *Cd86, Il1b, Cxcl2, and Chil3* in isolated immune cells from the myocardium of $Panx1^{fl/fl}$ and $Panx1^{MyHC6}$ mice treated with isoproterenol for 14 days (15 mg/kg/day). (N=4)

Data represented as mean $\pm$ SEM. Significance determined using Student's t-test (E).
